# CRCM5484: A BET- BDII Selective Compound With Differential Anti-Leukemic Drug Modulation

**DOI:** 10.1101/2021.12.13.472388

**Authors:** Kendall Carrasco, Camille Montersino, Carine Derviaux, Magali Saez-Ayala, Laurent Hoffer, Audrey Restouin, Rémy Castellano, Justine Casassa, Philippe Roche, Eddy Pasquier, Sébastien Combes, Xavier Morelli, Yves Collette, Stéphane Betzi

## Abstract

Differentially screening the Fr-PPIChem chemical library on the BET BRD4-BDII versus -BDI bromodomains led to the discovery of a BDII selective tetrahydropyridothienopyrimidinone (THPTP)-based compound. Structure-activity relationship (SAR) and hit-to-lead approaches allowed us to develop CRCM5484, a potent inhibitor of BET proteins with a preferential and 475-fold selectivity for the second bromodomain of the BRD3 protein (BRD3-BDII) over its first bromodomain (BRD3-BDI). Its very low activity was demonstrated in various cell-based assays, corresponding with recent data describing other selective BDII compounds. However, screening on a drug sensitivity and resistance-profiling platform revealed its ability to modulate the anti-leukemic activity in combination with various FDA-approved and/or in-development drugs in a cell- and context-dependent differential manner. Altogether, the results confirm the originality of the THPTP molecular mode of action in the BD cavity and its potential as starting scaffold for the development of potent and selective bromodomain inhibitors.

## INTRODUCTION

The bromodomain and extra-terminal (BET) family is composed of four epigenetic reader proteins, ubiquitous BRD4, BRD3 and BRD2 and testis-restricted BRDT. Each protein of this family includes a tandem small protein domain called bromodomain (BD) labeled BDI and BDII, both localized at the N-terminus of the proteins, and this domain can recognize and bind acetylated lysine on the histone tails. When BDs are bound to chromatin, they recruit transcription factors and play a role in gene transcription^1^. Pan-BET inhibitors are small compounds that can bind the eight BET BDs with similar potencies. This class of BD inhibitors has already been extensively studied for their therapeutic potential in both inflammation^2–8^ and oncology applications^9–14^. Among these well-studied compounds are JQ1^10,15^, OTX-015^16^ and I-BET-151^17^. Some of these molecules are currently being evaluated in clinical trials with oncology indications, confirming the interest for this protein family as therapeutic targets^18–20^. Nevertheless, toxicities and resistance phenomena have also been reported^19,21–23^. One of the main strategies to understand and overcome these issues has been to elucidate the functional contribution of the individual BDI and BDII domains in the BET family with the purpose of improving the adverse event profile observed in clinical trials.

Despite a high homology across the BDs in the BET family, several selective inhibitors of the BDI or BDII BET domains have recently emerged in the literature^24–27^, but few are currently engaged in clinical trials. The first developed selective compounds were BDI selective, the most extensively studied are olinone^28^, MS402^29^, and GSK778^30^. Developing BDII selective compounds was far more challenging, and only a handful of BDII selective inhibitors have been developed to date (Figure 1). The oldest compound, RVX-208 based on a quinazolinone chemical core, exhibited a selectivity of 20-fold with K_D_ values of 4100 nM and 200 nM (as measured by isothermal titration calorimetry (ITC)) against BRD4-BDI and BRD4-BDII, respectively^14,31^. GSK340 with a tetrahydroquinoxaline core is another BDII selective compound that exhibited K_D_ values of 3000 nM and 63 nM against BRD4-BDI and BRD4-BDII, respectively^32^ (as determined by BROMOScan). Very recently, ABBV-744 (tetrahydropyrrolopyridinone) was reported as a nanomolar BDII selective inhibitor and clinical candidate with a K_D_ measured by TR-FRET at 520 nM and 1.6 nM against BRD4-BDI and BRD4-BDII, respectively^33,34^. Finally, GSK046 (fluorobenzamide) and GSK620 (pyridinone), the latest additions to this list of BDII selective compounds, displayed IC_50_ values against the BDs of BRD3 that were 3600 nM and 98 nM for GSK046, and 2500 nM and 72 nM for GSK620, as determined by TR-FRET^30,35^. These compounds were evaluated for application in inflammatory disease, and the authors concluded that inhibition of the BET BDII domain may be a useful therapeutic strategy to counter inflammatory damage^30^.

**Figure 1.**
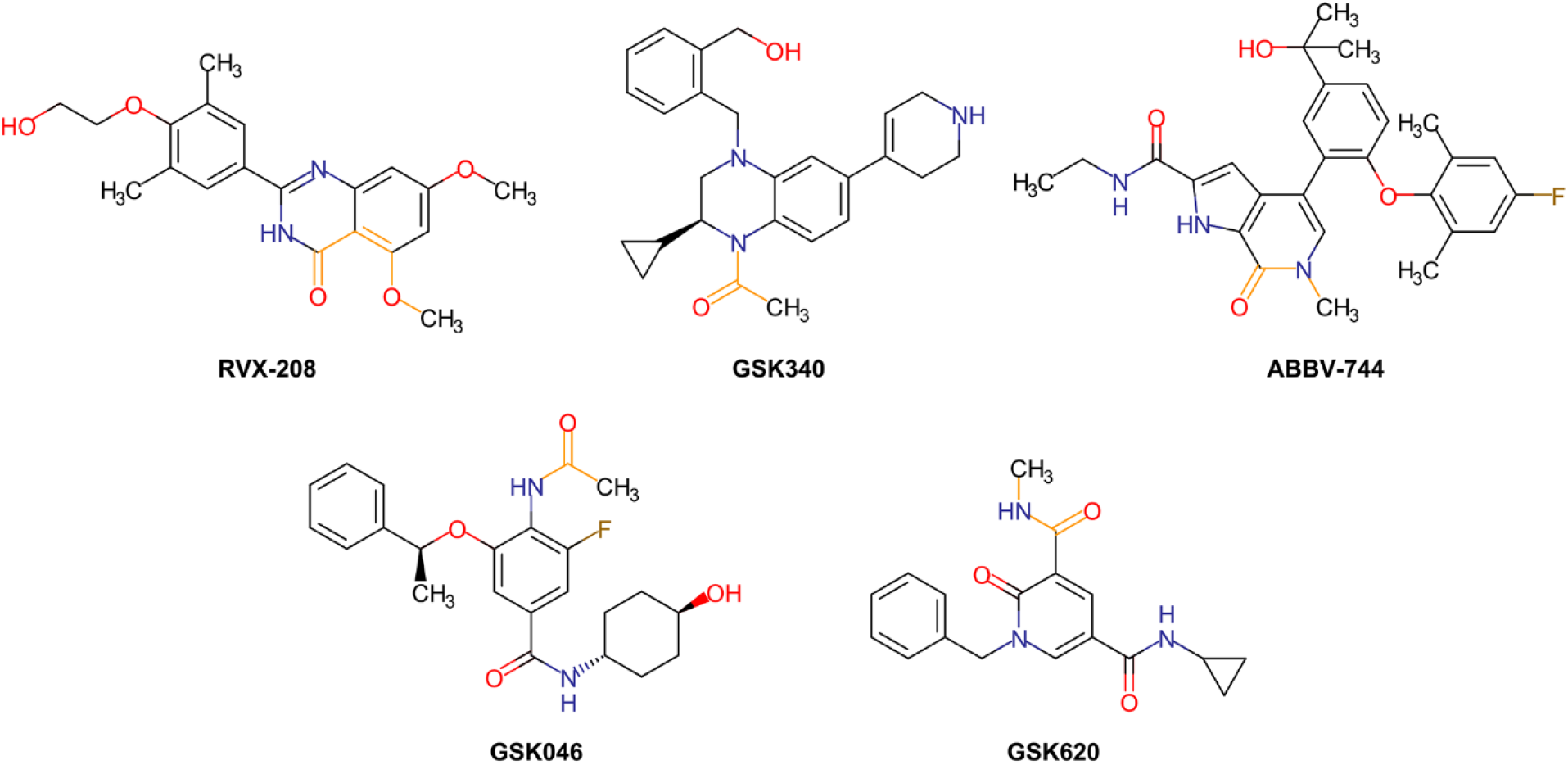
Chemical structures of the literature BDII selective inhibitors RVX-208, GSK340, ABBV-744, GSK046 and GSK620. Chemical moieties mimicking the histone acetylated lysine interaction with the key asparagine amino acid of the BDs are highlighted in orange.

In order to seek for new potential cores selective of either BDI or BDII domains, we miniaturized and robotized a 1536 wells format BD inhibitors assay and initiated a differential (BDI vs BDII) high-throughput screen (HTS) of the Fr-PPIChem library. Fr-PPIChem is a unique chemical library composed of 10314 diverse small molecules dedicated to protein-protein interaction (PPI) inhibition that displays considerably higher hit rates against PPIs than that of classical libraries^36^. Another fundamental advantage of this library is that its compounds as well as a series of structural analogs are commercially available. Moreover, medicinal chemistry filters were applied to remove undesirable structures (such as Pan-Assay Interference Compounds [PAINS], frequent hitters and toxic compounds) and to improve drug-likeness properties. From the Fr-PPIChem screen, an original tetrahydropyridothienopyrimidinone (THPTP)-based compound was identified as a bromodomain binder with a BDII selective profile that was further optimized through a structure activity relationship (SAR) campaign assisted by X-ray structural data. Our optimized hit - CRCM5484 - exhibits similar *in vitro* and in cellular activity to the most selective compounds described in the literature. Finally, a drug sensitivity response profile (DSRP) in a combination screen of CRCM5484 with 78 pharmacological agents pinpointed that CRCM5484 could potentiate the antiproliferative activity of certain drugs in patient-derived leukemia cells, which could be exploited for future developments. This potentiating effect is dependent upon the cellular context, and the drug response could be further potentiated with the addition of culture medium conditioned by human bone marrow stromal cells to mime the microenvironment of tumor cells; this highlights the importance of considering the impact of the tumor microenvironment when evaluating drug sensitivity in oncology.

## RESULTS

### THPTP core was identified by a “Fr-PPIChem” screening as a candidate for selective BDII BET inhibition

To develop our BDII selective BET inhibitor, we performed a differential screening on the “Fr-PPIChem” chemical library^36^. We used homogeneous time-resolved fluorescence (HTRF)^37^ to screen the 10314 compound library against both BDs of BRD4 (BRD4-BDI and BRD4-BDII). The scatter plot in Figure 2A confirms that the hits in the eight 1536-well screening plates were randomly distributed, thus experimental bias could be excluded. In addition, we measured several statistical data quality features for the screening. All these indicators were in agreement with industry standards, including Z’ factors of 0.83 and 0.87 and signal-to-noise ratios of 12.5 and 7.8 for BRD4-BDI and BRD4-BDII, respectively. The variability coefficient was also found to be below 5% for the whole screening. The screen designed to highlight BDII selective compounds identified 17 molecules with >30% BRD4-BDII inhibition and <30% BRD4-BDI screening at a final concentration of 1.25 μM. Among the molecules that met these criteria, 7 were selected for further evaluation (Table 1 and Figure S1 and S2A). The same selection threshold used to select BDI selective compounds would have led to 132 molecules, highlighting the challenge of developing BDII over BDI selective inhibitors.

**Figure 2.**
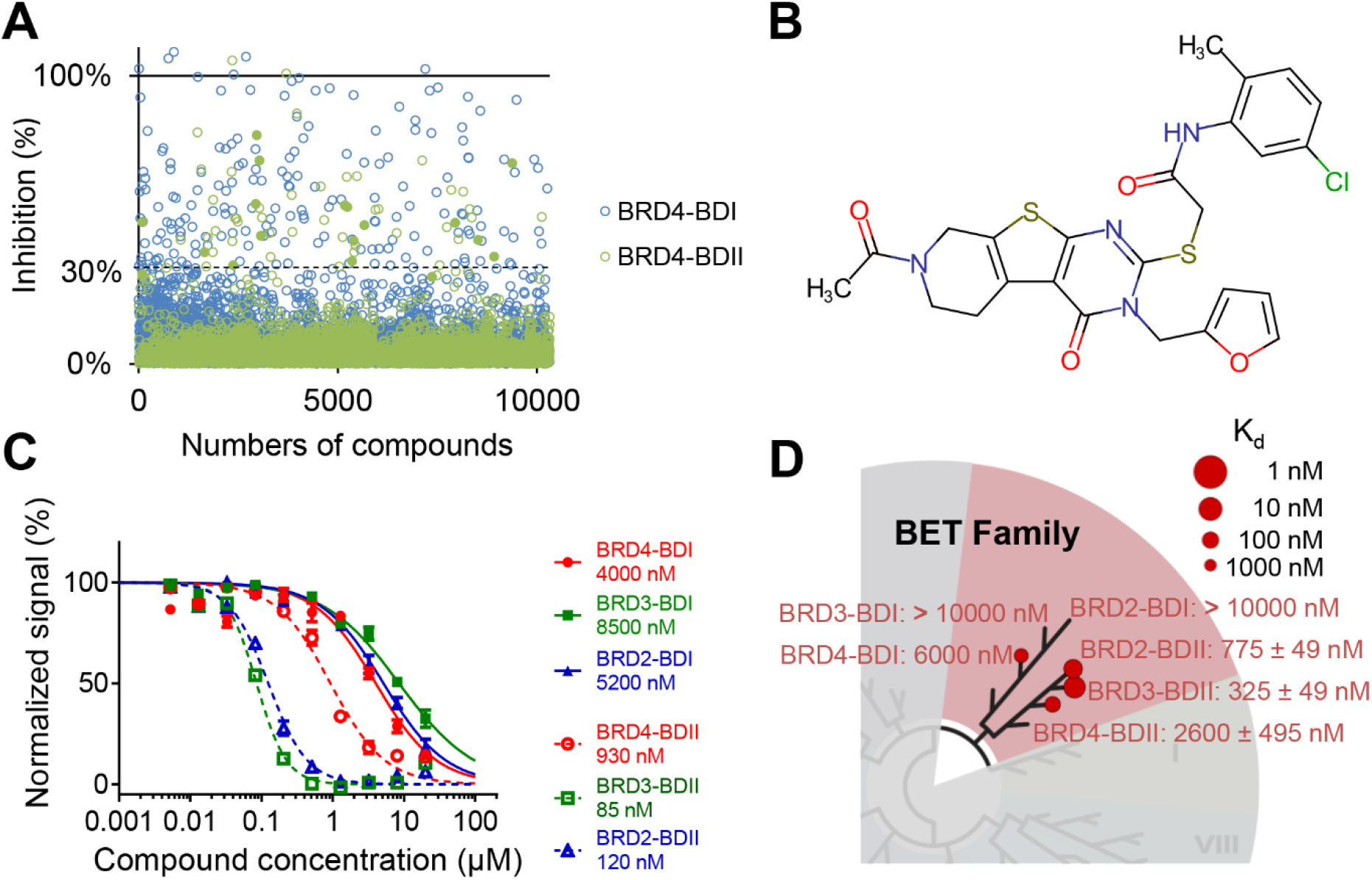
Fr-PPIChem screening and compound 1 characterization. (A) Scatter plot of Fr-PPIChem screening results against BRD4-BDI (empty blue circles) and BRD4-BDII (empty green circles). BDII selective compounds identified above the 30% threshold are shown as solid green circles. (B) Compound 1 chemical structure. (C) HTRF selectivity assay on the BDI (solid lines) and BDII (dashed lines) of BRD2 (blue), BRD3 (green) and BRD4 (red). (D) BROMOscan data on the BET bromodomain family for both BDs of BRD2, BRD3 and BRD4.

**Table 1.**
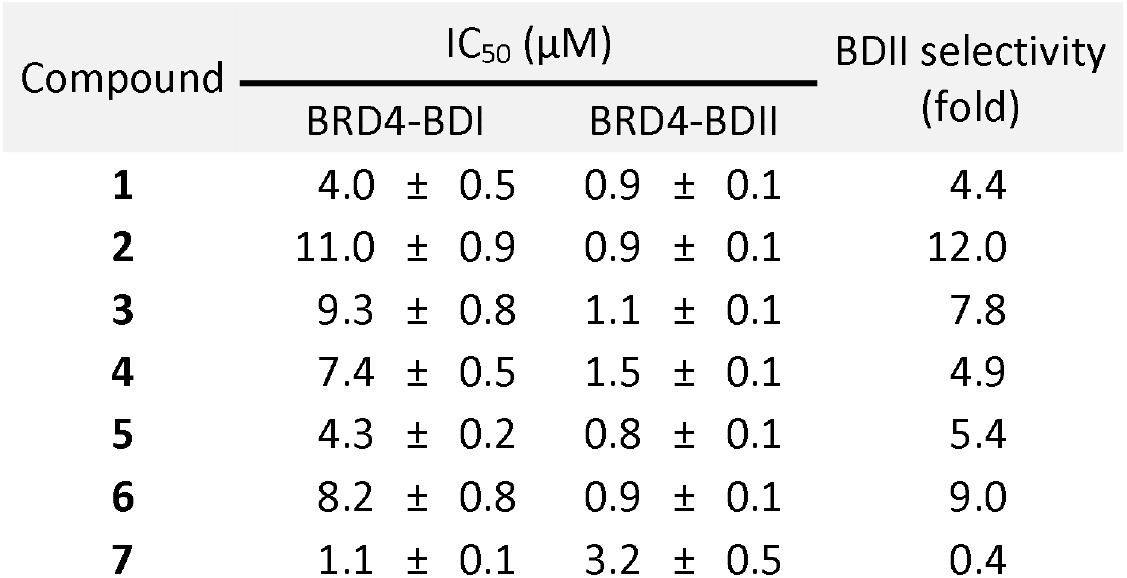
Screening results of seven BDII selective hits identified. The IC_50_ were obtained by HTRF assay on BRD4-BDI and BRD4-BDII.

We focused on compound 1 (Figure 2B) because it had a high affinity for BRD4-BDII (~900 nM) and because of its original THPTP chemical scaffold in the BET inhibitor chemical landscape. Indeed, only one related compound could be identified for this chemical scaffold in the literature as a BD binder in one crystal structure deposited in the PDB in 2016 (PDB ID: 4ZW1), but there was no follow-up study or publication exploiting these data (Figure S3A). The other identified compounds, a dimethyl isoxazole core of **2, 3** and **4,** has already been described in the BD inhibitor I-BET-151^25^ or GSK778^30^, and the trimethoxyphenyl core of 5 is found in DC-BD-03^38^ (Figure S2B). These results demonstrate the potential of the Fr-PPIChem chemical library to identify a new chemical core during a screening campaign.

The affinity and the selectivity of 1 were confirmed by two experiments: an HTRF selectivity assay on the six BDs of the BET family and a BROMOscan assay on the whole BD family tree (DiscoverX - Eurofins) (Figure 2C and D). These results confirm that 1 preferentially binds the BDII domains of BRD3 and BRD2, as confirmed by the IC_50_ values of 43 nM and 120 nM, respectively, which were measured by HTRF. The affinity for BRD4-BDII is significantly lower, with an IC_50_ of 930 nM. This was further supported by the BROMOscan results, in which the dissociation constants for the BDIIs of BRD4, BRD3 and BRD2 were measured at 2600 nM, 325 nM and 775 nM, respectively. This characteristic is common to other already-published BDII selective compounds, such as GSK046 and GSK620 that also bind preferentially the BDII sub-domains of BRD2 and BRD3 over BRD4-BDII^30^.

### THPTP optimization by SAR

To investigate the molecular mode of action of 1 and how its different chemical functions affect the affinity and selectivity toward BET BDs, we initiated an SAR campaign by ordering similar compounds from commercial libraries (SAR by catalog). The first five compounds purchased were selected because of the common 7-alcanoyl-5,6,7,8-tetrahydropyrido[4’,3’:4,5]thieno[2,3-d]pyrimidin-4(1H)-one core, as shown in Table 2. After measuring their affinities for both BDs of BRD3 by HTRF, we concluded that the *N*-acetylated pyrido moiety is a key binding component. This observation suggests there is a different binding mode than that of the 4ZW1 PDB compound in which the interaction is driven by the xylenol core (Figure S3).

**Table 2.** Structure Activity Relationship (SAR) by catalog. The IC_50_ were obtained by HTRF assay on BRD3-BDI and BRD3-BDII.

| Compound | R | Ar | $IC_{50}$ (nM) | | BDII selectivity (fold) |
| --- | --- | --- | --- | --- | --- |
|  |  |  | BRD3-BDI | BRD3-BDII |  |
| **1** | 2-furfuryl | 5-chloro-2-methylphenyl | 8500 ± 130 | 85 ± 3 | 100 |
| **8** | phenyl | 5-chloro-2-methylphenyl | > 25000 ± 1487 | 6300 ± 220 | > 4 |
| **9** | tetrahydrofuran-2-ylmethyl | 5-chloro-2-methylphenyl | 7800 ± 400 | 410 ± 10 | 19 |
| **10** | cyclohexyl | - | > 25000 ± 2435 | NA | NA |
| **11** | phenyl | phenyl | > 25000 ± 1726 | 5500 ± 3638 | > 5 |
| **12** | 2-furfuryl | phenyl | 14000 ± 510 | 140 ± 10 | 100 |

The presence of a furfuryl group as an R substituent also appears to be important for the selectivity profile. This substitution is indeed the only difference from 1, and the affinity for BRD3-BDII is degraded with IC_50_ values of 85 nM, 410 nM and 6300 nM for **1, 8** and **9,** respectively. Compound **12** is the best compound in this series, exhibiting an IC_50_ of 140 nM on BRD3-BDII and a selectivity of 100-fold. We hypothesized that the furan moiety adopts a position in close proximity to histidine H433 of BRD2-BDII, one of the main hot-spot amino acids that is targeted when developing BDII selective drugs. H433 is indeed one of the few residues discriminating BDII from BDI in an otherwise identical cavity. The only difference with 1 is the unsubstituted phenyl ring (Table 2). This observation suggests that aryl is important for binding to BD. The HTRF assay on BRD4, BRD3 and BRD2 for both BDs shows that **12** maintains the BDII selective profile (Figure S4A).

The importance of aryl prompted us to further investigate with another round of SAR by catalog, focusing on various phenyl substitutions. As the HTRF evaluation of the previous series identified solubility issues, we also included the predicted compound log P as an additional selection metric. We chose compounds with a log P below 4 because the predicted log P of 1 is 4.2 (Table3). The affinity and selectivity evaluation of this series by HTRF provided valuable SAR information. First, methyl, ethoxy or methoxy substituents in the -*Para* position degraded the affinity for BRD3-BDII, with IC_50_ values ranging from 570 to 880 nM against the 140 nM measured for **12.** This suggests the ring has a close proximity to the protein surface, causing steric hindrance when substituents are included. Substitution with a methoxy or ethoxy groups in the *-Ortho* position could increase the compound affinity because of additional hydrogen bonds. Compound **16,** with anisole, is the most interesting compound of this set, with affinities of 2 200 nM and 72 nM for BRD3-BDI and BRD3-BDII, respectively. The HTRF assay on both BDs of the BET family validated the BDII selective profile of **16** (Figure S4B).

**Table 3.** Structure Activity Relationship (SAR) of compound 12 benzyl ring. IC_50_ values were obtained by HTRF on BRD3-BDI and BRD3-BDII. Log P values were generated with Marvin Sketch.

| Compound | Ar | Log P | IC<sub>50</sub> (nM) |  | BDII selectivity (fold) |
| --- | --- | --- | --- | --- | --- |
|  |  |  | BRD3-BDI | BRD3-BDII |  |
| **1** | 5-Chloro-2-methylphenyl | 4.2 | 8500 ± 130 | 85 ± 3 | 100 |
| **12** | phenyl | 3.0 | 14000 ± 510 | 140 ± 10 | 100 |
| **13** | 3-Methoxyphenyl | 2.9 | 2800 ± 330 | 300 ± 12 | 9 |
| **14** | 2-Chlorophenyl | 3.7 | 2300 ± 180 | 100 ± 5 | 23 |
| **15** | 3-Methylphenyl | 3.6 | 5200 ± 2100 | 440 ± 34 | 12 |
| **16** | 2-Methoxyphenyl | 2.9 | 2200 ± 150 | 72 ± 3 | 31 |
| **17** | 4-Methoxyphenyl | 2.9 | 3600 ± 910 | 570 ± 22 | 6 |
| **18** | 4-Fluorophenyl | 3.2 | 3000 ± 2300 | 420 ± 18 | 7 |
| **19** | 2-Methylphenyl | 3.6 | 3200 ± 750 | 220 ± 7 | 15 |
| **20** | 3-Methylphenyl | 3.6 | 3100 ± 970 | 630 ± 23 | 5 |
| **21** | 3-Chlorophenyl | 3.7 | 4400 ± 1700 | 490 ± 15 | 9 |
| **22** | 2-Ethoxyphenyl | 3.2 | 3500 ± 730 | 150 ± 9 | 23 |
| **23** | 2,4-Difluorophenyl | 3.3 | 7500 ± 3300 | 380 ± 13 | 13 |
| **24** | 2-Fluorophenyl | 3.2 | 4200 ± 320 | 210 ± 6 | 20 |
| **25** | 3-Fluoro-4-methylphenyl | 3.7 | 6500 ± 5200 | 210 ± 6 | 31 |
| **26** | 3-Methylthiophenyl | 3.7 | 4500 ± 1900 | 410 ± 12 | 11 |
| **27** | 4-Chlorophenyl | 3.7 | 4500 ± 2000 | 530 ± 18 | 8 |
| **28** | 2,5-Difluorophenyl | 3.3 | 4100 ± 1600 | 170 ± 4 | 24 |
| **29** | 3-Acetylphenyl | 2.6 | 1500 ± 64 | 170 ± 6 | 9 |
| **30** | 4-Acetylphenyl | 2.6 | 1300 ± 45 | 320 ± 12 | 4 |
| **31** | 4-Ethoxyphenyl | 3.2 | 3800 ± 2500 | 880 ± 22 | 4 |
| **32** | 3-Chloro-4-fluorophenyl | 3.8 | 4900 ± 2700 | 740 ± 28 | 7 |
| **33** | 3,4-Difluorophenyl | 3.3 | 3800 ± 1500 | 470 ± 11 | 8 |

At this point, all purchasable analogs were evaluated, and we expanded the SAR study with in-house chemical synthesis. We pursued the optimization strategy by exploring additional modifications of the aromatic ring to increase the activity and solubility. With the addition of the picoline group, the predicted Log P decreased from 2.9 for compound 16 to 2.4 for compound 34 (CRCM5484). The 2-((7-acetyl-3-(furan-2-ylmethyl)-4-oxo-3,4,5,6,7,8-hexahydropyrido[4’,3’:4,5]thieno[2,3-d]pyrimidin-2-yl)thio)-*N*-(2-methylpyridin-3-yl)acetamide (CRCM5484, **34)** was prepared by convergent synthesis in three steps as depicted in scheme 1. A mixture of commercially available *N*-acetyl-4-piperidone, activated ethyl 2-cyanoacetate and elemental sulfur, was subjected to a *one-pot* Gewald reaction^39^ in presence of triethylamine as base to afford ethyl 6-acetyl-2-amino-4,5,6,7-tetrahydrothieno[2,3-c]pyridine-3-carboxylate (CRCM5484-2, **34-2),** in good yields. Reacting this latter with 2-furfuryl isothiocyanate in acetonitrile afforded corresponding 7-acetyl-3-(furan-2-ylmethyl)-2-mercapto-5,6,7,8-tetrahydropyrido[4’,3’:4,5]thieno[2,3-d]pyrimidin-4(3H)-one (CRCM5484-1, **34-1).** Finally, condensation with 2-chloro-*N*-(2-methylpyridin-3-yl)acetamide (CRCM5484-3, **34-3),** previously obtained by reaction of appropriate substituted aminopyridine with chloroacetyl chloride, afforded the expected THPTP (CRCM5484, **34)** in 90% yields.

**Scheme 1.**
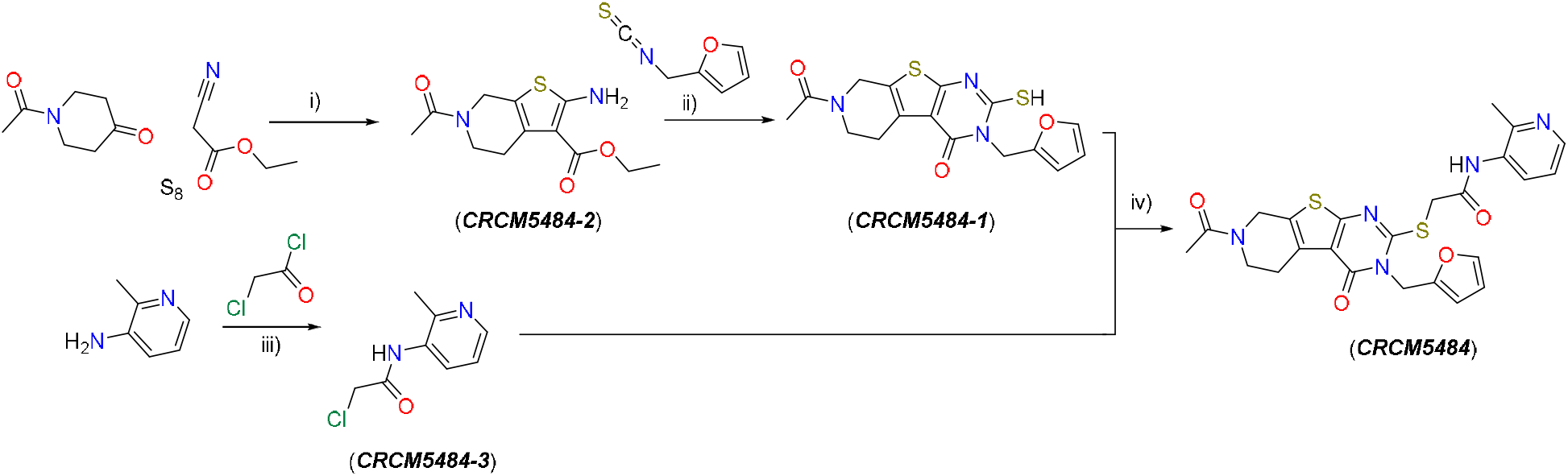
Reagents and Conditions: i) TEA, EtOH, reflux, 2h, 74%; ii) 2-furfurylisothiocyanate, K_2_CO_3_, MeCN, reflux, 12h, 10%; iii) chloroacetyl chloride, TEA, DCM, rt, 5h, 82%; iv) K_2_CO_3_, acetone, reflux, 2h, 90%.

### CRCM5484, a BDII selective BET inhibitor

CRCM5484 (Figure 3A) exhibited a BDII selective profile with IC_50_ values of 130 nM, 20 nM and 71 nM for the BDIIs and 1300 nM, 9500 nM and 4700 nM for the BDIs of BRD4, BDR3 and BRD2, respectively (Figure 3B). The affinity for BRD3-BDII was confirmed by isothermal titration calorimetry (ITC) with a 150 nM measured K_D_ (Figure 3C). The BROMOscan evaluation at 1μM confirmed the BDII selectivity profile (Figure 3D and Table S1) and also an overall BET family selectivity across the 32 evaluated domains. From this set of data, only two additional relatively lower affinity binders could be highlighted i.e. TRIM33 (52%) and CECR2 (54%) as compared to the 36, 37 and 41% inhibition measured for the BDII of BRD3, BRD2, and BRD4, respectively. A potential impact of these additional binders, either beneficial or detrimental, cannot be ruled out when assessing the biological effects of CRCM5484. The co-crystal structure of CRCM5484 in complex with BRD4-BDI and BRD2-BDII was obtained at 1.2 and 1.5 Å resolution, respectively (Figure 3E). As expected from the SAR study, the binding mode of CRCM5484 is different than that of the thienopyrimidine derivative compound in the 4ZW1 structure (Figure S3B). For CRCM5484, the N-acetyl from the tetrahydropyridinyl moiety is anchored in the binding pocket and mimics the acetylated lysine through direct hydrogen bonds with the key asparagine residues N140 and N429 in BDI and BDII, respectively. The acetamido group also stabilizes the water network at the bottom of the pocket. A major conformational change is also observed for CRCM5484 between the two BDs, as shown in Figure 3E and Figure S5. The picoline bound to BRD4-BDI (marked with an orange star Figure 3F) and the furan to BRD2-BDII (marked with a blue star Figure 3G) showed poor electronic density and their positioning were driven by geometry and residual electron density during refinement. In the BDII structure, the picoline moiety occupies the ZA channel and establishes vdW interactions with the WPF shelf residues W370, P371 and F372. This position is also stabilized by various interactions with water molecules and other protein residues. In BDI, this moiety is oriented toward the Y139 and N140 amino acids (Figure 3F and Figure S5A and E). The furyl moiety that we originally linked to the BDII selective profile does not establish direct interactions with H433, the main hot spot for the development of BDII selective drugs (Figure 3G and Figure S5C and F). The crystal structure links the BDII selectivity to a combination of direct or water-mediated anchor points that stabilize the compound in the cavity. In both structures, the furyl moiety binds a pocket formed by the WPF shelf and an aspartate residue (D145 for BRD4-BDI and D434 for BRD2-BDII). Altogether, these results confirm the originality of the THPTP molecular mode of action in the BD cavity and its potential as starting scaffold for the development of potent BD inhibitors.

**Figure 3.**
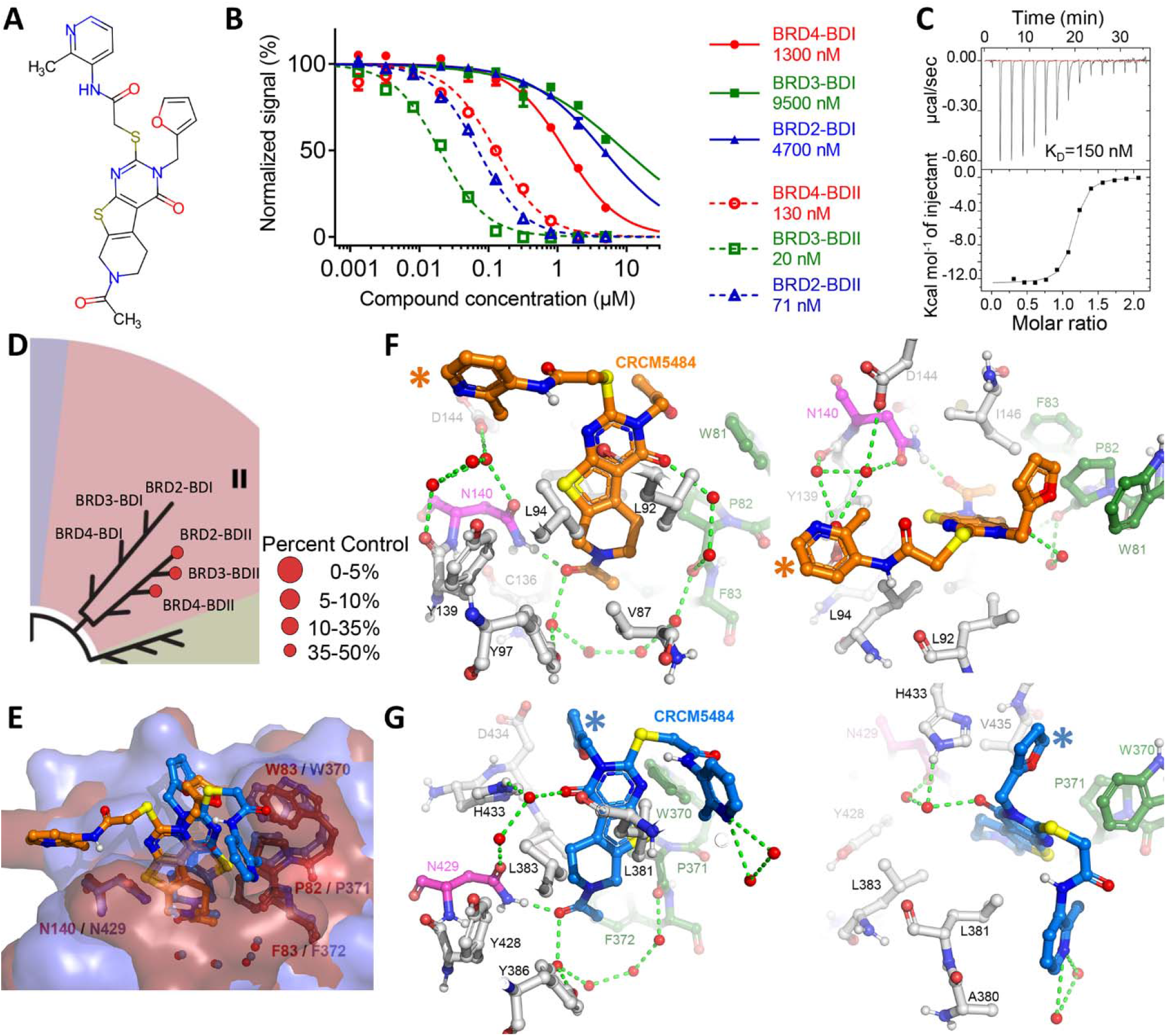
Characterization of CRCM5484. (A) Chemical structure. (B) HTRF selectivity assay on the BDI (solid lines) and BDII (dashed lines) of BRD2 (blue), BRD3 (green) and BRD4 (red). (C) Representative Isothermal Titration Calorimetry (ITC) experiment with CRCM5484 in the cell and BRD3-BDII in the syringe at 15°C. (D) BROMOscan data (DiscoverX) on the 8 BET bromodomains families. (E) Superposition of the co-crystal structures of CRCM5484 (orange)/BRD4-BDI (red) (PDB: 7Q3F) and CRM5484 (cyan) / BRD2-BDII (blue) (PDB: 7Q5O). Water molecules are shown as red and blue spheres for the BRD4-BDI and BRD2-BDII complexes, respectively. F and G) Co-Crystal structures of CRCM5484 with BRD4-BDI (Orange) and BRD2-BDII (Blue) showing the hydrogen bond interactions as green dotted lines, the canonical asparagine binding residue as magenta, the WPF shelf as green and the rest of the domain as white balls and sticks. The picoline (orange *) and furan moiety (blue *) showed poor electronic density and their positioning were driven by geometry and residual electron density during refinement.

### CRCM5484 has similar biological effects as other BDII selective compounds in cell-based assays

The biological effects of CRCM5484 were evaluated and compared to the reference BDII selective compounds RVX-208 and GSK046, and the pan-BET inhibitor OTX-015. First, treatment of the MOLM-14 leukemic cell line showed that CRCM5484, similar to RVX-208 and GSK046, exhibited >100x lower impact on cell growth and viability than that of OTX-015, with EC_50_ values of 8.5 μM, 6.7 μM, 14 μM and 0.08 μM, respectively (Figure 4A). Second, the transcriptional impact of these various BD inhibitors on *MYC* gene expression^40^ was compared by quantitative RT-PCR. All tested molecules induced a concentration-dependent decrease in *MYC* gene expression, with the pan-BET inhibitor displaying the highest potency (IC_50_ = 0.6 μM), and the BDII selective compounds were at least 5x less efficient with IC_50_ values ranging between 3 and 7 μM (CRCM5484) (Figure 4B). These evaluations were replicated in three additional leukemia cell lines, the known bromodomain inhibitor sensitive CCRF-CEM in addition to MOLM-14 and the two resistant cell lines, NB4 and K-562. Overall, CRCM5484 and the two other evaluated BDII selective compounds (RVX-208 and GSK046) displayed little to no cytotoxicity on the 4 cell lines with EC_50_ > 5μM, however MOLM-14 and CCRF-CEM cells appeared slightly more responsive to the drugs (Figure S6A). MYC gene expression profile was also similar for these cell lines (Figure S6B). As BDII inhibition was recently shown to modulate the inflammatory stimulation of gene transcription in MHC-I antigen presentation pathway gene components^25^, we also evaluated the impact of CRCM5484 on K-562 leukemic cells stimulated with the proinflammatory cytokine interferon-_⍰_ (IFN_⍰_). As recently reported, the pan-BET inhibitor totally prevented IFN_⍰_-induced expression of MHC-I while CRCM5484 partially inhibited this induction in a concentration-dependent manner, with a similar efficiency to GSK046 and in agreement with Gilan et al^30^ (Figure 4C). Although it cannot be excluded that the two molecules hold distinct cell penetration capabilities, these results show that CRCM5484 exhibits comparable activity to the compound GSK046 in these cellular assays.

**Figure 4.**
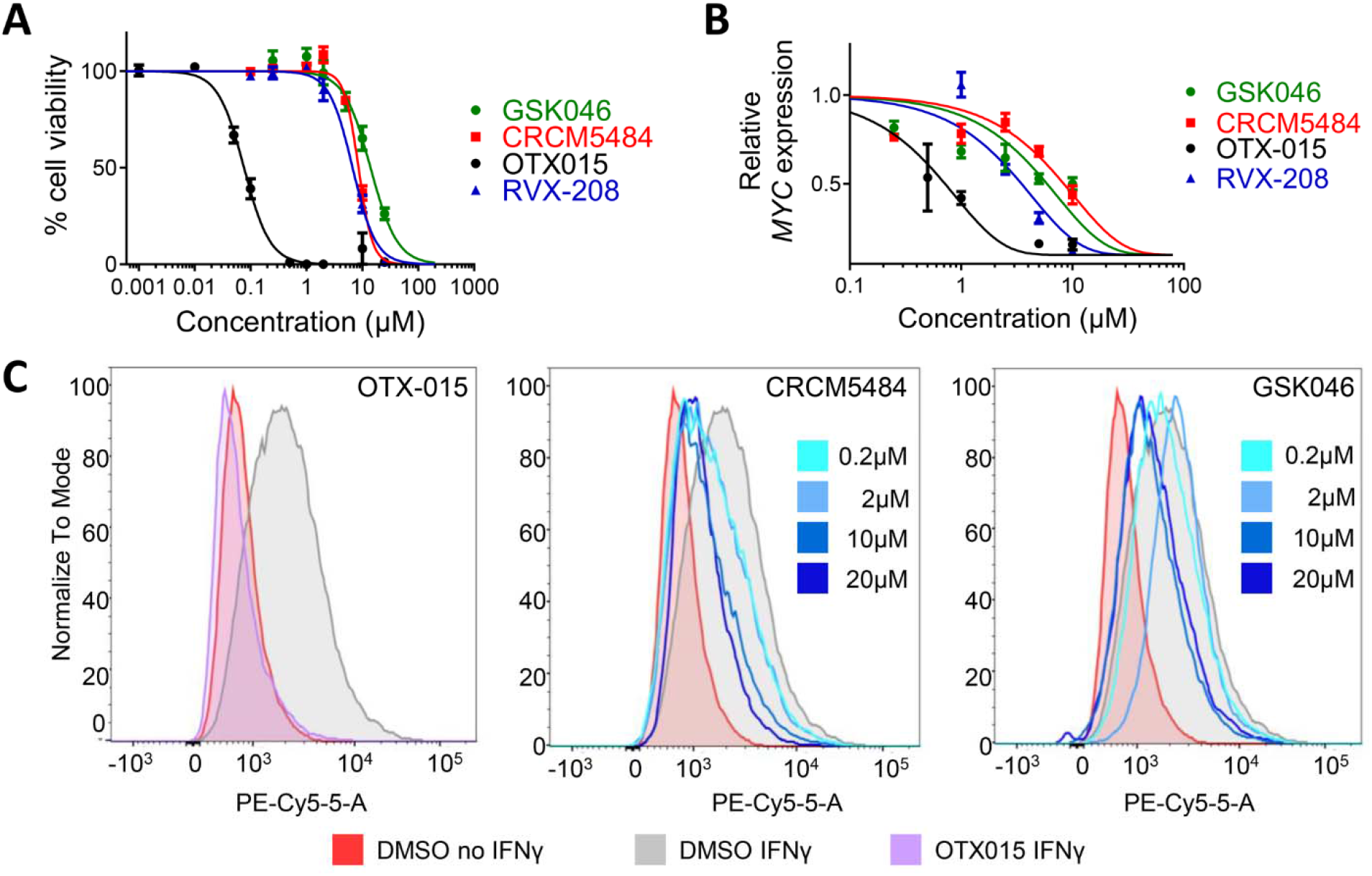
CRCM5484 compared to other BDII selective compounds. (A) Cell viability assay on MOLM-14 leukemic cells performed after 72h incubation with OTX015 (black), RVX-208 (blue), GSK046 (green) and CRCM5484 (red). (B) Quantification of *MYC* expression by RT-qPCR following 2h incubation with OTX015 (black), RVX-208 (blue), GSK046 (green) and CRCM5484 (red). (C) Flow cytometry analysis of MHC-I expression in K-562 cells following stimulation by IFN_⍰_ and treatment with DMSO, OTX015, GSK046 or CRCM5484.

### Cell- and context-dependent modulation of drug sensitivity of leukemic cells by CRCM5484

Because of the limited impact of BDII selective compounds as single agents in cell viability/growth assays, we next questioned the potential to use them in combination with other known compounds demonstrating anti-leukemic activity. Using our previously described DSRP platform^41^, CRCM5484 was added to 3 distinct leukemia cell lines (MOLM-14, U-937 and CCRF-CEM), as well as to primary patient-derived xenografted (PDX) acute myeloid leukemia cells at its 40% effective concentration (EC_40_) (previously established for each individual cell line and PDX-derived cell cultures, see Table S2 for details), in combination with a panel of 78 drugs. Since the biological impact of selective BDII inhibition is best evidenced in inflammatory conditions and because drug sensitivity and resistance profiles of primary leukemia cell cultures have been shown to be differentially modulated in the presence of bone-marrow-derived stromal cell conditioned medium (BMSC-CM)^42^, PDX-derived leukemia cells^43^ were cultured in both the absence and presence of BMSC-CM. The fold potentiation ratio was then calculated as the ratio of each tested drug EC_50_ value determined in the presence and absence of CRCM5484 (Figure 5A and B). By arbitrarily setting a threshold of potentiation of at least 3-fold, CRCM5484 potentiated the cell viability/growth inhibitory activity of 16 drugs: 10 in MOLM-14, 1 in U-937, 3 in the CCRF-CEM cell line and 2 in the PDX-derived leukemia cells in the absence of BMSC-CM. CRCM5484-potentiated drugs varied with the investigated cell cultures, since only 1 out of these 16 drugs, namely birinapant, was potentiated in at least 3 out of the 4 distinct cell cultures, including PDX-derived leukemia cells. Strikingly, adding BMSC-CM to the PDX-derived leukemia cell cultures greatly modified the drug sensitivity and resistance profiles; 7 additional drugs were potentiated >3-fold by CRCM5484 (Figure 5B), but none of these drugs were also potentiated in the established leukemia cell lines (Figure 5A).

**Figure 5.**
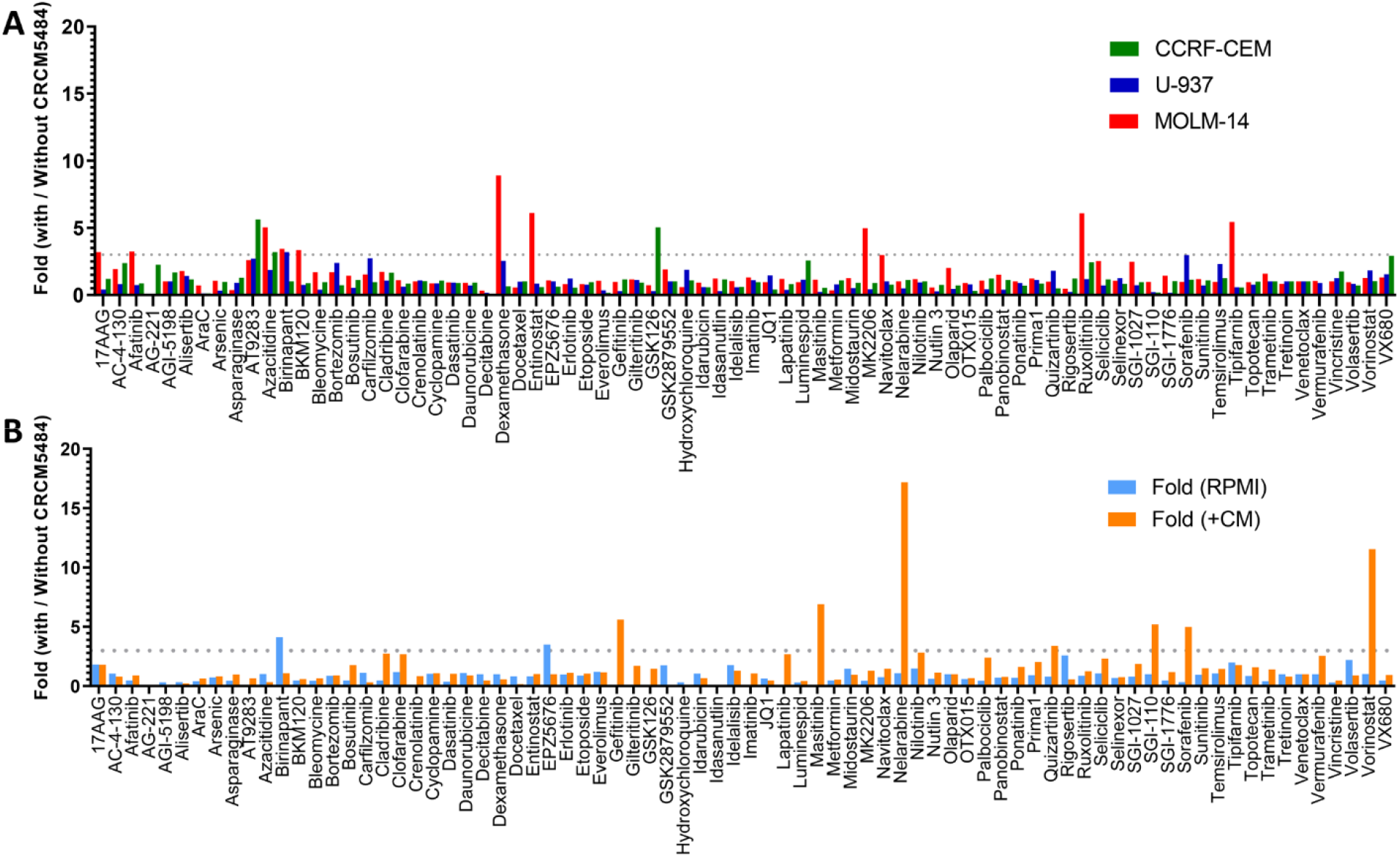
CRCM5484 potentiation of pharmacological agents. (A) Fold of drug cytotoxic effects with or without CRCM5484 at its EC_40_ (Table S2) in CCRF-CEM (green), U-937 (blue) and MOLM-14 (red) cell lines. (B) Fold of drug cytotoxic effects with or without CRCM5484 with or without addition of bone-marrow-derived stromal cell conditioned medium (BMSC-CM).

To confirm these results on selected drug combinations, we developed dose-response experiments with combinatorial effects analyses on PDX-derived leukemia cells. First, we focused on birinapant as the sole drug identified in our screening strategy in 3 out of 4 cell cultures, including PDX-derived leukemia cells cultured in the absence of BMSC-CM. As shown in figure 6A, a dose-dependent but limited potentiation of birinapant cell viability/growth inhibitory activity by CRCM5484 was observed, which was scored as mild additivity to low synergy by Combenefit^44^ matrix analysis (Figure 6B). Out of the 7 drugs in which the anti-leukemic activity was potentiated by CRCM5484 in the presence of BMSC-CM, we focused on vorinostat. Indeed, for four of these drugs (nelarabine, quizartinib, sorafenib and gefitinib), the potentiation by CRCM5484 most likely resulted from increased resistance induced by BMSC-CM, as previously reported^42^. Furthermore, among the remaining three drugs (vorinostat, masitinib and SG1-110), vorinostat displayed the highest potentiation ratio (11-fold) and has been previously described to synergize with pan-BET inhibitors^45^. Dose-response experiments with combinatorial effect analyses confirmed the absence of vorinostat potentiation by CRCM5484 in the absence of BMSC-CM (Figure 6C and D) and dose-dependent potentiation in the presence of BMSC-CM (Figure 6E and F), which was scored as synergistic by Combenefit matrix analysis at a range of vorinostat concentrations that are clinically achievable^46^.

**Figure 6.**
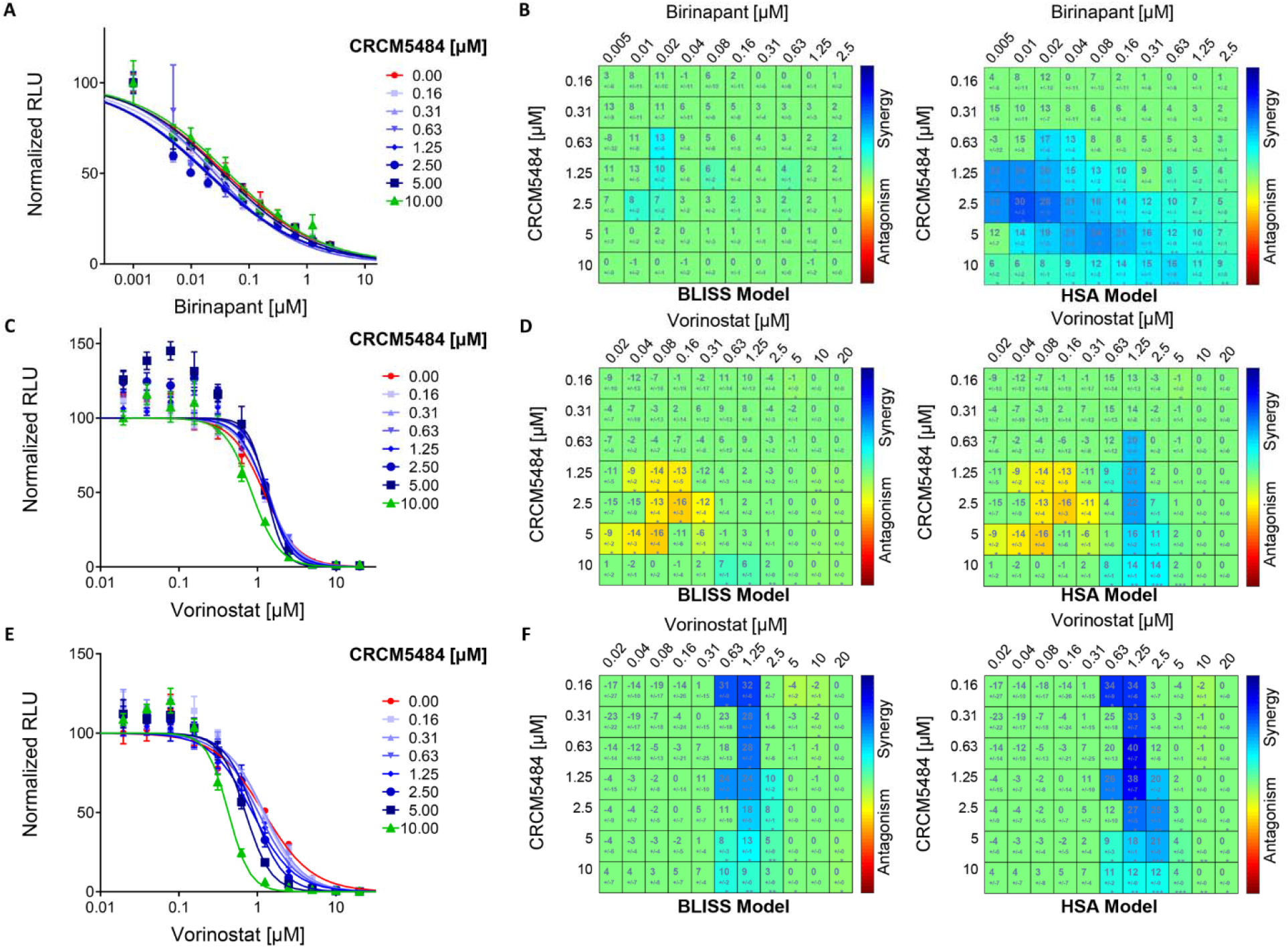
Validation of CRCM5484 potentiation of pharmacological agents in conditioned medium in the presence/absence of BMSC-CM. Dose response curves of birinapant (A) or vorinostat (C and E) cytotoxic effect at a concentration range of CRCM5484 in the absence (A and C) or the presence (E) of BMSC-CM. Combination effect matrix between birinapant (B) and vorinostat (D and F) in combination with CRCM5484 analyzed using the BLISS (right) and HSA model (left) in the absence (B and D) or the presence (F) of BMSC-CM. The synergy/antagonism score and its statistical significance is specified for each combination. Significant combinations are colored in an antagonism/synergy scale (red to blue) and stars indicate the level of significance.

Altogether, these results support the differential ability of CRCM5484 to potentiate anti-leukemic drug activity in a cell and culture condition-dependent manner.

## DISCUSSION AND CONCLUSIONS

The development of selective BDI and/or BDII compounds has been strongly motivated by the encouraging results of first-generation pan-BET inhibitors in various clinical trials, which have even led to remissions in some cancer patients^47–49^. However, the initial enthusiasm was rapidly tempered by the short response durations observed at tolerated doses and the description of resistance mechanisms that may contribute to these limitations. Therefore, developing more selective molecules was quickly proposed as well as evaluating combinations with standard treatments and/or other targeted therapies to reduce side effects while improving efficacy. Designing inhibitors with a selective profile could also lead to a better understanding of the role played by each domain in the BD family, which is currently under debate, and to propose new original chemistry-based molecules that could be developed as drug candidates.

In this study, we describe how a differential screen of BDI and BDII domains of the BET family against our in-house “Fr-PPIChem” chemical library led to the identification of a BD binder with a preference for BDII of the BRD3 and BRD2 proteins from the BET family. This compound has an original selectivity profile based on a THPTP core and has not been previously described as a BD binder in the literature. To optimize this screening hit, a series of five compounds was first purchased during SAR by a catalog campaign, and the molecular mode of action of the best compounds was investigated. The acetamido group appeared to be crucial for compound anchoring in the binding pocket while the furyl moiety was responsible for the selectivity profile.

In a second series of nineteen compounds, phenyl substitution could adjust the affinity of the compounds to BDs, and we observed that substitutions in the -Para position, such as ethoxy, methoxy, methyl, chloro or fluoro groups decreased compound affinity with an IC_50_ >500 nM on BRD3-BDII. We predicted that the best compounds of the series established additional hydrogen bonds with a hydrogen bond donor or acceptor in the -Ortho position on the phenyl ring. Within this series, compound **16** exhibited the best inhibitory potential with IC_50_ values measured at 2200 and 72 nM for the BDI and BDII of BRD3 respectively, and a 31-fold selectivity ratio.

A last step in the chemical synthesis aimed at improving the compound selectivity and solubility and led to CRCM5484, our lead compound that bears a picoline moiety instead of the initial aryl moiety. This compound is a BDII selective inhibitor that exhibits a similar activity in TR-FRET to GSK046 or RVX-208, the reference BDII selective compounds in the literature. Comparative analysis of the X-Ray structures of CRCM5484 in complex with BRD4-BDI and BRD2-BDII supported the observations from the SAR by catalog and revealed that the compound was rotated by 180° in the cavity. In BRD4-BDI, the picoline moiety established no interaction with the protein, while in BDII, the picoline moiety was stabilized by vdW contacts in the protein WPF shelf region.

We then evaluated the activity of our THPTP-based BDII inhibitor in a variety of cell-based assays, in comparison with two reference BDII selective inhibitors: the recently developed GSK046^30^, and RVX-208, the earliest described BDII compound^14^. Consistent with the activities reported for the latter compounds, when used as a single agent, CRCM5484 presented a weak impact on the viability/proliferation of the leukemic MOLM-14 cell line (as well as K562, CCRF-CEM and NB4) with EC_50_ values >100x compared to those of the pan-BET compound OTX-015, as well as a limited impact on the down-regulation of *MYC* gene transcription, a well-documented transcriptional target of BET inhibitors.

In parallel, the work of Gilan *et al* demonstrated the contributions of BDII to the transcriptional induction of genes, and BDII notably controlled the cell surface expression of the MHCI complex by inflammatory cytokines and its modulation by the BDII selective compound GSK046. Consistent with these observations, our results demonstrate that CRCM5484 modulates IFN_⍰_-induced MHCI expression in a manner comparable to that of GSK046 but is less efficient than the pan-BET inhibitor OTX-015.

Based on these observations, we then assessed the ability of CRCM5484 to potentiate the antileukemic activity of a panel of 78 drugs with marketing authorization and/or in clinical development. Our screening approach highlights several observations. First, when CRCM5484 is added in a single dose corresponding to its previously measured EC_40_ in each of the cell models evaluated, the antiproliferative activity of the tested drugs was potentiated up to 9-fold, depending on the cell line and including primary AML cells derived from a PDX. Second, the drugs potentiated by CRCM5484 varied greatly depending on the cell line evaluated and the presence of inflammatory conditioned medium from bone marrow stromal cells for the test performed on primary PDX-derived AML cells (up to 17-fold increased antiproliferative activity for the later). Third, the selection of the two drugs with the best combination potential confirmed the potentiation effect of CRCM5484 in dose-response experiments, which were performed on primary PDX-derived AML cells with additivity/low synergy effects in combination with birinapant and synergy effects in combination with vorinostat, in the presence of conditioned medium from bone marrow stromal cells.

To our knowledge, these results are the first illustration of a combination effect of BET inhibitors with birinapant, a second mitochondrial-derived activator of caspases (SMAC) mimetic, in which the rationale for the combination effect remains to be unveiled and better documented. It should be noted that although this potentiation was observed for two of the three cell lines as well as the primary AML cells from our PDX model, it was not observed for the latter in the presence of BMSC-CM.

The potentiation effect of the histone deacetylase inhibitor vorinostat by pan-BET inhibitors has recently been reported in cutaneous T-cell lymphoma^50^. However, it is important to note that this effect was only observed for primary AML cells from our PDX model and in the presence of BMSC-CM. It therefore seems important to confirm these observations and in particular to extend them to a panel of primary AML samples.

Nevertheless, our results provide motivation to further explore the capacities of CRCM5484 and, more broadly, selective BDII inhibitors to potentiate the anti-leukemic activity of FDA-approved drugs or molecules in development; and the inhibitors should be examined in terms of the potentially reduced side effects due to their better selectivity in various preclinical cancer models.

## EXPERIMENTAL SECTION

### Chemistry

For chemical synthesis, all compounds are >95% pure by HPLC analysis. The Table S3 summarizes all compounds information (SMILES, CAS, provider, and provider ID). Quality evaluations as provided by the company (NMR/HPLC) for all purchased compounds are included as supplementary files. Commercially available reagents and solvents were used without further additional purification. Thin layer chromatography (TLC) was performed on precoated aluminum sheets of silica (60 F254 nm, Merck) and visualized using short-wave UV light. Reaction monitoring and purity of synthesized compounds were recorded by using analytical Agilent Infinity high performance liquid chromatography (HPLC) with DAD at 254 nM column Agilent Poroshell 120 EC-C18 2.7μm (4.6 x 50 mm), mobile phase (A: 0.1% FA H2O, B: 0.1% FA MeCN), flow rate 0.3 mL/min, time/%B 0/10, 4/90, 7/90, 9/10, 10/10. Column chromatography was performed on a Reveleris purification system using Reveleris Flash silica cartridges. Petroleum refers to the fraction with distillation range 40-65 °C. ^1^H and ^13^C NMR spectra were recorded by using a Bruker AC 400 spectrometer. Chemical shifts, (δ) are reported in ppm and coupling values (J) in hertz. Abbreviations for peaks are, br: broad, s: singlet, d: doublet, t: triplet, q: quadruplet, quint: quintuplet, sext: sextuplet, sept: septuplet and m: multiplet). The spectra recorded are consistent with the proposed structures. Low-resolution mass spectra were obtained with Agilent SQ G6120B mass spectrometer in positive and negative electrospray mode.

### Ethyl 6-acetyl-2-amino-4,5,6,7-tetrahydrothieno[2,3-c]pyridine-3-carboxylate (CRCM5484-2, 34-2)^51^

To a suspension of ethyl 2-cyanoacetate (5.66 g, 50 mmol), *N*-acetyl-4-piperidinone (5.71 g, 50 mmol), triethylamine (7 mL, 50 mmol) and sulfur (1.60 g, 50 mmol) in ethanol (100 mL) was heated at 70 °C for 2 h. The resulting mixture was concentrated under reduced pressure and the residue was successively diluted with DCM (100 mL), washed with H_2_O (2 x 30 mL), brine (30 mL), and dried over Na_2_SO_4_. The solvent was distillated off under reduced pressure and the crude product was purified by crystallization from Et_2_O to afford ethyl 6-acetyl-2-amino-4,5,6,7-tetrahydrothieno[2,3-c]pyridine-3-carboxylate **CRCM5484-2** (10 g, 75%), as a yellow powder. NMR (400 MHz, CDCl_3_) δ 4.71 (brs, 2H), 4.52-4.37 (m, 2H), 4.26 (q, *J* = 7.2 Hz, 2H), 3.79-3.65 (m, 2H), 2.86 (brs, 2H), 2.17 (brs, 3H), 1.36 (t, *J* = 7.2 Hz, 3H); LCMS C_12_H_16_N_2_O_3_S Rt = 6.908 min, ESI+ m/z = 269.1 (M+H).

### 7-Acetyl-3-(furan-2-ylmethyl)-2-mercapto-5,6,7,8-tetrahydropyrido[4’,3’:4,5]thieno[2,3-d]pyrimidin-4(3H)-one (CRCM5484-1, 34-1)

To a solution of **CRCM5484-2** (5.36 g, 20 mmol) and 2-furfuryl isothiocyanate (2.76 g, 20 mmol) in acetonitrile (50 ml), was added anhydrous potassium carbonate (2.76 g, 20 mmol). The resulting mixture was heated under reflux overnight. After cooling down at room temperature, the precipitate formed was filtered off, and resuspended, under stirring, in 2N HCl aqueous solution (30 mL) for 15 min. The solid was collected by filtration to afford the 7-acetyl-3-(furan-2-ylmethyl)-2-mercapto-5,6,7,8-tetrahydropyrido[4’,3’:4,5]thieno[2,3-d]pyrimidin-4(3H)-one **CRCM5484-1** (700 mg, 10%), as a light yellow powder. NMR (400 MHz, DMSO) δ 13.71 (brs, 1H), 7.54 (d, *J* = 1.2 Hz, 1H), 6.37 (dd, *J* = 3.1, 1.2 Hz, 1H), 6.31 (d, *J* = 3.1 Hz, 1H), 5.57 (s, 2H), 4.64 (s, 0.5H), 4.59 (s, 1.5H), 3.71-3.67 (m, 2H), 2.94 (t, *J* = 5.4 Hz, 1.5H), 2.81 (t, *J* = 5.4 Hz, 0.5H), 2.10 (s, 2.25H), 2.05 (s, 0.75H); LCMS C_16_H_15_N_3_O_3_S_2_ Rt = 7.145 min, ESI+ m/z = 362.1 (M+H).

### 2-Chloro-*N*-(2-methylpyridin-3-yl)acetamide (CRCM5484-3, 34-3)^52^

At room temperature, to a solution of 3-amino-2-methylpyridine (2.16 g, 20 mmol) in dichloromethane (20 mL), were successively added chloroacetyl chloride (1.6 mL, 20 mmol) and triethylamine (2.79 mL, 20 mmol). The resulting mixture was stirred for 5 h, and concentrated under reduced pressure. The residue was purified by column chromatography, eluent DCM-MeOH (80:20) to afford the 2-chloro-*N*-(2-methylpyridin-3-yl)acetamide **34-3.** (3.02 g, 82%), as a white powder. LCMS C_8_H_9_ClN_2_O Rt = 2.179 min, ESI+ m/z = 185.1 (M+H).

### 2-((7-Acetyl-3-(furan-2-ylmethyl)-4-oxo-3,4,5,6,7,8-hexahydropyrido[4’,3’:4,5]thieno[2,3-d]pyrimidin-2-yl)thio)-*N*-(2-methylpyridin-3-yl)acetamide (CRCM5484, 34)

To a suspension of **CRCM5484-1** (181 mg, 0.5 mmol) and anhydrous potassium carbonate (170 mg, 1.23 mmol) in dry acetone (15 mL) was added **CRCM5484-3** (139 mg, 0.75 mmol). The resulting mixture was refluxed for 1 h and concentrated under reduced pressure. The residue was purified by column chromatography, eluent DCM-MeOH (95:5) to afford the 2-((7-acetyl-3-(furan-2-ylmethyl)-4-oxo-3,4,5,6,7,8-hexahydropyrido[4’,3’:4,5]thieno[2,3-d]pyrimidin-2-yl)thio)-*N*-(2-methylpyridin-3-yl)acetamide **CRCM5484** (230 mg, 90%), as a white powder. CRCM5484 purity was analyzed by HPLC as shown in Figure S7A. ^1^H NMR (400 MHz, CDCl_3_) δ 8.71 (brs, 1H), 8.36-8.31 (m, 1H), 8.27 (d, *J* = 5.1 Hz, 1H), 7.35 (brs, 1H), 7.22 (dd, *J* = 8.1, 5.1 Hz, 1H), 6.45 (d, *J* = 3.2 Hz, 1H), 6.33 (dd, *J* = 3.2, 1.9 Hz, 1H), 5.35 (s, 2H), 4.77 (s, 1.5H), 4.64 (s, 0.5H), 4.11 (s, 2H), 3.88 (t, *J* = 5.6 Hz, 0.5H), 3.74 (t, *J* = 5.6 Hz, 1.5H), 3.16 (t, *J* = 5.6 Hz, 1.5H), 3.10 (t, *J* = 5.6 Hz, 0.5H), 2.43 (s, 3H), 2.21 (s, 2.25H), 2.19 (s, 0.75H); ^13^C NMR (10 MHz, CDCl_3_) δ 169.53, 169.40, 166.59, 166.53, 161.69, 157.39, 157.21, 157.06, 156.86, 149.48, 149.34, 147.70, 145.29, 142.81, 132.14, 131.48, 130.29, 130.22, 129.63, 128.77, 126.97, 121.76, 118.72, 118.57, 110.65, 110.51, 110.43, 45.23, 43.51, 41.13, 40.53, 38.68, 36.44, 26.14, 25.26, 22.03, 21.42, 20.90; LCMS C_24_H_23_N_5_O_4_S_2_ Rt = 6.298 min, ESI+ m/z = 510.1 (M+H) as shown in Figure S7B, >98% pure.

### Protein expression and purification

For homogeneous time resolved fluorescence (HTRF) experiments, all 6 BDs of BRD4, BRD3 and BRD2 (3 BDI and 3 BDII domains) synthetic genes that include a Tobacco Etch Virus (TEV) cleavage site were purchased from LifeTechnology in a pDONR transport vector before cloning into a pDEST™15 expression vector for GST affinity purification. Protein production and purification was carried out using similar protocols and buffers used for the His-BRD4-BDI system. Purification was carried on GST affinity resin (Thermo Scientific) and reduced glutathione was used for protein release. GST-BRD4-BDI was further purified by size exclusion chromatography on a Superdex 16/60 Hiload column (GE Healthcare) using 20 mM TRIS pH=8.0, 150 mM NaCl Buffer.

For isothermal titration calorimetry (ITC), BRD3-BDII was produced and purified using a histidine tag affinity chromatography from a pDEST™17 expression vector containing a TEV protease cleavage site. After size exclusion chromatography, fractions presenting pure BRD3-BDII were pooled and concentrated to 7 mg/mL in 10 mM HEPES pH=7.5, 150 mM NaCl, 1mM DTT buffer. A final step of buffer exchange was performed using a PD10 column from GE healthcare prior to the experiment to remove the DTT.

For X-Ray crystallography, BRD4-BDI was produced and purified using a histidine tag affinity chromatography as described by Filipakopoulos et al^49^. For these experiments, a pNIC28-BSA4 expression vector containing BRD4-BDI and a TEV protease cleavage site has been kindly provided by Stefan Knapp laboratory from the SGC at the University of Oxford. After size exclusion chromatography, fractions of pure BRD4-BDI after TEV cleavage of the histidine tag were pooled and concentrated to 25 mg/mL.

For X-Ray crystallography, BRD2-BDII was produced from a pDEST™17 expression vector containing a TEV protease cleavage site and purified using a histidine tag affinity chromatography. After size exclusion chromatography, fractions of pure BRD2-BDII after TEV cleavage were pooled and concentrated to 20 mg/mL.

### Homogeneous Time-Resolved Fluorescence

HTRF assays were performed in white 1536 Well Small Volume™ HiBase Polystyrene Microplates (Greiner) with a total working volume of 4 μL using the conditions described in Table S4. 5 nL of compounds were dispensed (2 x 2.5 nL fixed dispensing using an Echo550 (Labcyte)) from a concentration stock of 1 mM (100% DMSO) for the Fr-PPIchem primary screening after addition of 15 nL DMSO per well to reach the final 1.25μM concentration used in the assay (0.5% final DMSO). Dose response evaluations of commercial and synthesized compounds were performed similarly from 10 mM stock solutions (100% DMSO). For the screening, the inhibition percentage was calculated and compared to the reference compound, JQ1. The IC_50_ measurements were carried out in triplicates. All HTRF reagents were purchased from CisBio Bioassays and reconstituted according to the supplier protocols. HTRF signals were measured, after a final incubation (6h at room temperature), using a PHERAstar FS (BMG Labtech) with an excitation filter at 337 nm and fluorescence wavelength measurement at 620 and 665 nm, using an integration delay of 60 μs and an integration time of 500 μs. Results were analyzed with a two-wavelengths signal ratio: [intensity (665 nm)/intensity (620 nm)]*10^4^. Percentage of inhibition was calculated using the following equation: % inhibition = [(compound signal) - (min signal)] / [(max signal) - (min signal)] * 100, where ‘max signal’ is the signal ratio with the compound vehicle alone (DMSO) and ‘min signal’ the signal ratio without peptide. For IC_50_ measurements, values were normalized and fitted with Prism (GraphPad) using the following equation: Y = 100 / (1 + ((X / IC_50_)^Hill slope)).

### BROMOscan

BROMOscan bromodomain profiling was provided by Eurofin DiscoverX Corp. Determination of the K_D_ between test compounds and DNA tagged bromodomains was achieved through binding competition against a proprietary reference immobilized ligand. CRCM5484 was used at 1μM.

### Isothermal Titration Calorimetry

ITC was used to evaluate the thermodynamics parameters of the binding between BRD4-BDI, BRD2-BDI and BRD3-BD2 and CRCM5484, using ITC conditions previously described by Filippakopoulos et al^53^. Purified proteins were extensively dialyzed in the ITC buffer containing 10mM Hepes pH=7.5 and 150 mM NaCl. Compound was diluted directly in the last protein dialysate prior to experiments. Titrations were carried out on a MicroCal ITC200 microcalorimeter (GE Healthcare, Piscataway, NJ). Each experiment was designed using a titrant concentration (protein in the syringe) set 10 to 15 times the analyte concentration (compound in the cell generally between 10 and 35 μM) and using 13 injections at 15 °C. A first small injection (generally 0.2 μL) was included in the titration protocol to remove air bubbles trapped in the syringe prior titration and/or take into account syringe predilution in the cell during equilibration. Raw data were scaled after setting the zero to the titration saturation heat value. Integrated raw ITC data were fitted to a one site non-linear least squares fit model using the MicroCal Origin plugin as implemented in Origin 7 (Origin Lab). Finally, ΔG and TΔS values were calculated from the fitted ΔH and KA values using the equations ΔG = -R.T.lnKA and ΔG = ΔH - TΔS.

### X-Ray crystallography

BRD4-BDI/CRCM5484 and BRD2-BDII/CRCM5484 co-crystallizations were performed at respectively, 4°C (277 K) and 12 °C (285 K) using the hanging drop vapor diffusion method. For the BRD4-BDI/CRCM5484, a solution of 25 mg/mL of BRD4-BDI with 1mM of compound was mixed at a 1:1 ratio with the precipitant solution (300 mM NaNO_3_, 16% PEG3350, 5% Ethylene glycol) and crystals grew to diffracting quality within 3-5 days.

For the BRD2-BDII/CRCM5484 complex, 20 mg/mL of BRD2-BDII and 2.5 mM of compound CRCM5484 preparation was mixed at a 1:1 ratio with the precipitant solution (25% PEG3350, 100 mM potassium thiocyanate) and crystals grew to diffracting quality within 10 days. Crystals grew to diffracting quality within 1 month.

Crystals were cryo-protected using the precipitant solution supplemented with 10% ethylene glycol and were flash frozen in liquid nitrogen. Data were collected at the ESRF beamlines ID30A-1 and at the SOLEIL beamline Proxima1. Indexing, integration, and scaling were performed using XDS. Initial phases were calculated by molecular replacement with Phaser MR (CCP4 suite) using a model of the first domain of BRD4 (extracted from the Protein Data Bank accession code: 2OSS); for BRD2-BDII, using a model extracted from the Protein Data Bank accession code: 5XHK. Initial models for the protein and the ligands were built in COOT. The cycles of refinement were carried out with phenix.refine (Phenix v1.13). Data collection and refinement statistics can be found in Table S5. The models and structures factors have been deposited with Protein Data Bank accession code for BRD4-BDI in complex with compound CRCM5484: 7Q3F, and BRD2-BDII in complex with the compound CRCM5484: 7Q5O.

### Cell culture

The human leukemia cell lines U-937, CCRF-CEM and K-562 were cultured in RPMI 1640 supplemented with 10% FBS. MOLM-14 were cultured in MEMalpha supplemented with 10% FBS. NB4 were cultured in Dulbecco’s Modified Eagle Medium supplemented with 10% FBS. All cell lines were incubated at 37°C with 5% CO2, were diluted every 2 to 3 days and maintained in an exponential growth phase. These cell lines have been routinely tested for mycoplasma contamination.

### Cell viability assay

MOLM-14, NB4, CCRF-CEM and K562 were seeded at 4000, 5000, 8000 and 6000 cells/well in 96-well plates, respectively. After 24h, cells were treated with a range of concentrations of CRCM5484, OTX015, RVX208 or GSK046 and after 72h drug incubation, metabolic activity was detected by addition of Alamar blue and spectrophotometric analysis using a PHERAstar plate reader (BMG labtech). Cell viability was determined and expressed as a percentage of untreated control cells. The determination of IC_50_ values was performed by using the following equation: Y=100/(1+((X/IC_50_)^Hillslope)).

### *MYC* gene expression analysis by RT-qPCR

MOLM-14 cells were treated with a range of concentrations of CRCM5484, OTX-015, RVX-208 or GSK046 for 2h. NB4, CCRF-CEM and K-562 cells were treated with 1 μM of OTX-015, 5 μM of CRCM5484, 5 μM of GSK046 or 5 μM of RVX-208 for 2h. RNA extraction was performed using RNeasy kit (Qiagen). OneScript^®^ Plus cDNA Synthesis Kit (abm) was used to prepare cDNA from purified RNA. Real-time qPCR was performed using SsoAdvanced Universal SYBR^®^ Green Supermix (Bio-Rad) and a CFX384™ Real-Time System device (Bio-Rad). Analysis performed on duplicate PCR data per biological replicate. Four biological replicates were used for DMSO, CRCM5484, GSK046, RVX208 and OTX015 treated cells. *MYC* gene expression level was determined using the ΔΔCt method, normalized to *GAPDH* control gene, using pre-designed KiCqStart™ primers (H_MYC_1, H_GAPDH_1; Merck).

#### Inhibition of MHC expression

The MHC-I expression was carried out by fluorescence-activated cell sorting analysis using antibodies against HLA-A/B/C (HLA-ABC Monoclonal Antibody (W6/32), PE-Cyanine5, eBioscience) in K-562 cells after stimulation with IFN-γ (10 ng/ml, PHC4031 life technologies) and treatment with DMSO, OTX-015, CRCM5484, or GSK046 for 48 hours. Following incubation and washing, samples were analyzed on a LSR II flow cytometer (Becton Dickinson) using flowjo V-10.8.1 software.

### Drug sensitivity and resistance assays

The drug screening library included 78 substances consisting of conventional chemotherapeutics and a broad range of targeted oncology compounds. Each drug was plated on 96-well plates at 4 concentrations covering a 1000-fold concentration range. 10-20000 cells were seeded per well in 96-well plates and incubated in the presence of compounds (4 concentrations generated after a 1 log serial dilution from the starting concentration defined in Table S2 for each drug/cell line) in presence or absence of CRCM5484 in a humidified environment at 37°C and 5% CO_2_. After 72h (cell lines) - 144h (PDX) of treatment, cell viability was measured using the CellTiter-Glo Luminescent Cell Viability Assay as described by the manufacturer (Promega Corporation). The Luminescence was measured using a Centro luminometer LB960 (Berthold). Effective half-maximal concentration values (EC_50_) were deduced from dose-response curves obtained using GraphPad Prism 6 software (GraphPad Software, Inc.). Primary AML cells were collected after two passages in mice from a previously described Patient-Derived Xenograft mouse model^43^ and were cultured with RPMI 20% FBS, 50% bone-marrow-derived stromal cell conditioned medium (HS-5 Cell line) or RPMI 20% FBS, as described^39^.

### Synergy assays

Median effect analysis was performed through Combenefit software^44^ based on Chou and Talalay method using growth inhibition obtained from CellTiter-Glo luminescent assay. The combination effect was compared to the cytotoxicity of each drug alone. We performed combination analyses using Bliss and HSA methods. The drug interactions were specified using Matrix Synergy Plot. The synergy assays were carried out in triplicates

## Supporting information

Supplementary_Informations

Providers_Data

experimental-data

## ASSOCIATED CONTENT

### Supporting Information

See figures S1 to S7 and Tables S1 to S5:

Figure 1: HTRF evaluation of the FR-PPIChem screening hits, Figure 2: chemical structures, Figure 3: binding mode comparison between CRCM5484 and 4ZW1, Figure 4: selectivity evaluation of compound **12** and compound **16,** Table 1: BROMOscan evaluation of CRCM5484, Figure 5: binding mode study of CRCM5484, Table 2: maximal concentration used for the 78 evaluated compounds in the drug sensitivity and resistance profiling; Figure 6: CRCM5484 HPLC analysis; Table 3: Compounds SMILES, Provider, Provider ID and CAS number; Table 4: HTRF experimental conditions, Table 5: X-ray data collection and refinement statistics. Quality evaluations as provided by the company (NMR/HPLC) are included as supplementary files (file Carrasco-et-al-Providers.zip) for all purchased compounds. CSV file listing compounds formula strings with their associated biochemical and biological data (Carrasco-et-al-Data.csv).

## AUTHOR INFORMATION

### Author Contributions

K.C. and C.D. carried out the small molecules screen; K.C. and S.B. performed the biochemical and biophysical evaluation; Chemical synthesis was performed by S.C. and J.C.; P.R. and L.H. performed the chemoinformatics and molecular modeling analysis; C.M., K.C., E.P. and A.R. carried out the cellular evaluations. The manuscript was written through contributions of all authors. S.B., Y.C. and X.M. supervised and conceived the project. All authors have given approval to the final version of the manuscript.

## PDB ID CODES

PDB codes for BRD4-BDI and BRD2-BDII with bound CRCM5484 are 7Q3F and 7Q5O. Authors will release the atomic coordinates upon article publication.

## ACKNOWLEDGMENTS

This study was supported by research funding from the “Agence Nationale de la Recherche” (ANR-15-CE18-0023). This study was also partly supported by research funding from the “Canceropôle PACA”, “Institut National du Cancer” and “Région Provence-Alpes-Côte d’Azur” (“Prematuration Grant” 2017 attributed to X.M. & “Emergence Grant” 2017 attributed to E.P) and from the “Fondation ARC” (PJA 20171206125 grant attributed to X.M.), K.C. was supported by a fellowship from the “Fondation ARC”. We would like to thank the FRISBI ANR-10-INSB-05-01 grant for access time on the structural-biology platform of the AFMB laboratory and Dr. Gerlind Sulzenbacher for the “block allocation group” management. We acknowledge the European Synchrotron Radiation Facility and the SOLEIL synchrotron for provision of synchrotron radiation facilities and the staff at beamline ID30A-1, Proximal and Proxima2. We thank the HiTS, TrGET and flow cytometry platforms of the CRCM and their staff. We thank Dr Etienne Rebuffet from iSCB team for fruitful discussions about our structural data.

## ABBREVIATIONS

BET: Bromodomain and Extra-Terminal
BD: Bromodomain
PPI: Protein-Protein Interactions
ITC: Isothermal Titration Calorimetry
PAINS: Pan-Assay Interference Compounds
HTS: High-Throughput Screen
THPTP: tetrahydropyridothienopyrimidinone
SAR: Structure Activity Relationship
DSRP: Drug Sensitivity Response Profile
HTRF: Homogeneous Time-Resolved Fluorescence
IFN_⍰_: Pro-Inflammatory Cytokine Interferon-_⍰;_
BMSC-CM: Bone-Marrow-Derived Stromal Cell Conditioned Medium
HPLC: High Performance Liquid Chromatography

## GRAPHIC FOR MANUSCRIPT

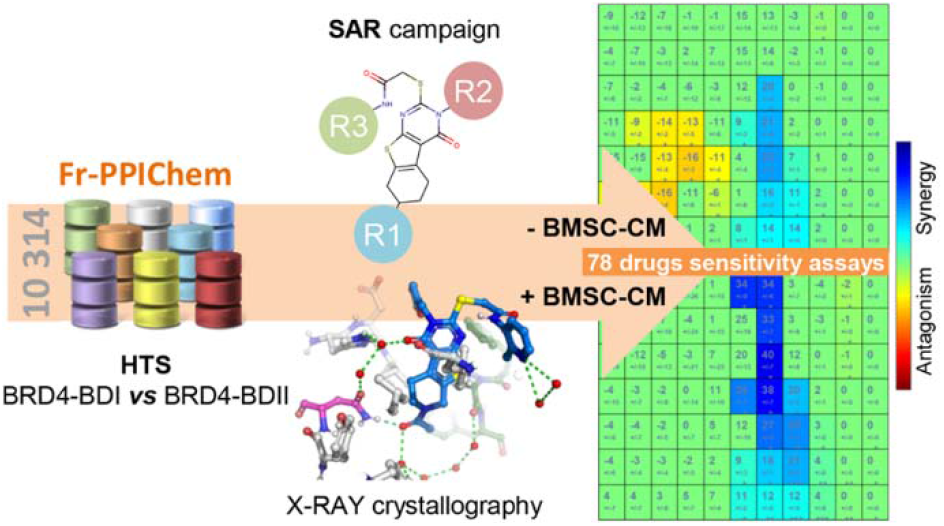

## REFERENCES

(1) Fujisawa, T.; Filippakopoulos, P. Functions of Bromodomain-Containing Proteins and Their Roles in Homeostasis and Cancer. Nat Rev Mol Cell Biol 2017, 18 (4), 246–262.

(2) Tough, D. F.; Tak, P. P.; Tarakhovsky, A.; Prinjha, R. K. Epigenetic Drug Discovery: Breaking through the Immune Barrier. Nat Rev Drug Discov 2016, 15 (12), 835–853.

(3) Tough, D. F.; Prinjha, R. K. Immune Disease-Associated Variants in Gene Enhancers Point to BET Epigenetic Mechanisms for Therapeutic Intervention. Epigenomics 2017, 9 (4), 573–584.

(4) Bandukwala, H. S.; Gagnon, J.; Togher, S.; Greenbaum, J. A.; Lamperti, E. D.; Parr, N. J.; Molesworth, A. M. H.; Smithers, N.; Lee, K.; Witherington, J.; Tough, D. F.; Prinjha, R. K.; Peters, B.; Rao, A. Selective Inhibition of CD4+ T-Cell Cytokine Production and Autoimmunity by BET Protein and c-Myc Inhibitors. PNAS 2012, 109 (36), 14532–14537.

(5) Schilderink, R.; Bell, M.; Reginato, E.; Patten, C.; Rioja, -->Inmaculada; Hilbers, F. W.; Kabala, P. A.; Reedquist, K. A.; Tough, D. F.; Tak, P. P.; Prinjha, R. K.; de Jonge, W. J. BET Bromodomain Inhibition Reduces Maturation and Enhances Tolerogenic Properties of Human and Mouse Dendritic Cells. Molecular Immunology 2016, 79, 66–76.

(6) Belkina, A. C.; Nikolajczyk, B. S.; Denis, G. V. BET Protein Function Is Required for Inflammation: Brd2 Genetic Disruption and BET Inhibitor JQ1 Impair Mouse Macrophage Inflammatory Responses. The Journal of Immunology 2013, 190 (7), 3670–3678.

(7) Nicodeme, E.; Jeffrey, K. L.; Schaefer, U.; Beinke, S.; Dewell, S.; Chung, C.; Chandwani, R.; Marazzi, I.; Wilson, P.; Coste, H.; White, J.; Kirilovsky, J.; Rice, C. M.; Lora, J. M.; Prinjha, R. K.; Lee, K.; Tarakhovsky, A. Suppression of Inflammation by a Synthetic Histone Mimic. Nature 2010, 468 (7327), 1119–1123.

(8) Zhang, Q.; Qian, J.; Zhu, Y. Targeting Bromodomain-Containing Protein 4 (BRD4) Benefits Rheumatoid Arthritis. Immunology Letters 2015, 166 (2), 103–108.

(9) Prinjha, R. K.; Witherington, J.; Lee, K. Place Your BETs: The Therapeutic Potential of Bromodomains. Trends in Pharmacological Sciences 2012, 33 (3), 146–153.

(10) Filippakopoulos, P.; Qi, J.; Picaud, S.; Shen, Y.; Smith, W. B.; Fedorov, O.; Morse, E. M.; Keates, T.; Hickman, T. T.; Felletar, I.; Philpott, M.; Munro, S.; McKeown, M. R.; Wang, Y.; Christie, A. L.; West, N.; Cameron, M. J.; Schwartz, B.; Heightman, T. D.; La Thangue, N.; French, C. A.; Wiest, O.; Kung, A. L.; Knapp, S.; Bradner, J. E. Selective Inhibition of BET Bromodomains. Nature 2010, 468 (7327), 1067–1073.

(11) Chaidos, A.; Caputo, V.; Gouvedenou, K.; Liu, B.; Marigo, I.; Chaudhry, M. S.; Rotolo, A.; Tough, D. F.; Smithers, N. N.; Bassil, A. K.; Chapman, T. D.; Harker, N. R.; Barbash, O.; Tummino, P.; Al-Mahdi, N.; Haynes, A. C.; Cutler, L.; Le, B.; Rahemtulla, A.; Roberts, I.; Kleijnen, M.; Witherington, J. J.; Parr, N. J.; Prinjha, R. K.; Karadimitris, A. Potent Antimyeloma Activity of the Novel Bromodomain Inhibitors I-BET151 and I-BET762. Blood 2014, 123 (5), 697–705.

(12) da Motta, L. L.; Ledaki, I.; Purshouse, K.; Haider, S.; De Bastiani, M. A.; Baban, D.; Morotti, M.; Steers, G.; Wigfield, S.; Bridges, E.; Li, J.-L.; Knapp, S.; Ebner, D.; Klamt, F.; Harris, A. L.; McIntyre, A. The BET Inhibitor JQ1 Selectively Impairs Tumour Response to Hypoxia and Downregulates CA9 and Angiogenesis in Triple Negative Breast Cancer. Oncogene 2017, 36 (1), 122–132.

(13) Rhyasen, G. W.; Hattersley, M. M.; Yao, Y.; Dulak, A.; Wang, W.; Petteruti, P.; Dale, I. L.; Boiko, S.; Cheung, T.; Zhang, J.; Wen, S.; Castriotta, L.; Lawson, D.; Collins, M.; Bao, L.; Ahdesmaki, M. J.; Walker, G.; O’Connor, G.; Yeh, T. C.; Rabow, A. A.; Dry, J. R.; Reimer, C.; Lyne, P.; Mills, G. B.; Fawell, S. E.; Waring, M. J.; Zinda, M.; Clark, E.; Chen, H. AZD5153: A Novel Bivalent BET Bromodomain Inhibitor Highly Active against Hematologic Malignancies. Mol Cancer Ther 2016, 15 (11), 2563–2574.

(14) Picaud, S.; Wells, C.; Felletar, I.; Brotherton, D.; Martin, S.; Savitsky, P.; Diez-Dacal, B.; Philpott, M.; Bountra, C.; Lingard, H.; Fedorov, O.; Müller, S.; Brennan, P. E.; Knapp, S.; Filippakopoulos, P. RVX-208, an Inhibitor of BET Transcriptional Regulators with Selectivity for the Second Bromodomain. PNAS 2013, 110 (49), 19754–19759.

(15) Filippakopoulos, P.; Picaud, S.; Fedorov, O.; Keller, M.; Wrobel, M.; Morgenstern, O.; Bracher, F.; Knapp, S. Benzodiazepines and Benzotriazepines as Protein Interaction Inhibitors Targeting Bromodomains of the BET Family. Bioorg Med Chem 2012, 20 (6), 1878–1886.

(16) Noel, J. K.; Iwata, K.; Ooike, S.; Sugahara, K.; Nakamura, H.; Daibata, M. Abstract C244: Development of the BET Bromodomain Inhibitor OTX015. Mol Cancer Ther 2013, 12 (11 Supplement), C244–C244.

(17) Seal, J.; Lamotte, Y.; Donche, F.; Bouillot, A.; Mirguet, O.; Gellibert, F.; Nicodeme, E.; Krysa, G.; Kirilovsky, J.; Beinke, S.; McCleary, S.; Rioja, I.; Bamborough, P.; Chung, C.-W.; Gordon, L.; Lewis, T.; Walker, A. L.; Cutler, L.; Lugo, D.; Wilson, D. M.; Witherington, J.; Lee, K.; Prinjha, R. K. Identification of a Novel Series of BET Family Bromodomain Inhibitors: Binding Mode and Profile of I-BET151 (GSK1210151A). Bioorg. Med. Chem. Lett. 2012, 22 (8), 2968–2972..

(18) Liu, Z.; Wang, P.; Chen, H.; Wold, E. A.; Tian, B.; Brasier, A. R.; Zhou, J. Drug Discovery Targeting Bromodomain-Containing Protein *4*. J. Med. Chem. 2017, 60 (11), 4533–4558.

(19) Doroshow, D. B.; Eder, J. P.; LoRusso, P. M. BET Inhibitors: A Novel Epigenetic Approach. Ann Oncol 2017, 28 (8), 1776–1787.

(20) Alqahtani, A.; Choucair, K.; Ashraf, M.; Hammouda, D. M.; Alloghbi, A.; Khan, T.; Senzer, N.; Nemunaitis, J. Bromodomain and Extra-Terminal Motif Inhibitors: A Review of Preclinical and Clinical Advances in Cancer Therapy. Future Science OA 2019, 5(3) FSO372.

(21) Andrieu, G.; Belkina, A. C.; Denis, G. V. Clinical Trials for BET Inhibitors Run Ahead of the Science. Drug Discovery Today: Technologies 2016, 19, 45–50.

(22) Postel-Vinay, S.; Herbschleb, K.; Massard, C.; Woodcock, V.; Soria, J.-C.; Walter, A. O.; Ewerton, F.; Poelman, M.; Benson, N.; Ocker, M.; Wilkinson, G.; Middleton, M. First-in-Human Phase I Study of the Bromodomain and Extraterminal Motif Inhibitor BAY 1238097: Emerging Pharmacokinetic/Pharmacodynamic Relationship and Early Termination Due to Unexpected Toxicity. European Journal of Cancer 2019, 109, 103–110.

(23) Odenike, O.; Wolff, J. E.; Borthakur, G.; Aldoss, I. T.; Rizzieri, D.; Prebet, T.; Hu, B.; Dinh, M.; Chen, X.; Modi, D.; Freise, K. J.; Jonas, B. A. Results from the First-in-Human Study of Mivebresib (ABBV-075), a Pan-Inhibitor of Bromodomain and Extra Terminal Proteins, in Patients with Relapsed/Refractory Acute Myeloid Leukemia. JCO 2019, 37 (15_suppl), 7030–7030.

(24) Rianjongdee, F.; Atkinson, S. J.; Chung, C.-W.; Grandi, P.; Gray, J. R. J.; Kaushansky, L. J.; Medeiros, P.; Messenger, C.; Phillipou, A.; Preston, A.; Prinjha, R. K.; Rioja, I.; Satz, A. L.; Taylor, S.; Wall, I. D.; Watson, R. J.; Yao, G.; Demont, E. H. Discovery of a Highly Selective BET BD2 Inhibitor from a DNA-Encoded Library Technology Screening Hit. J Med Chem 2021, 64 (15), 10806–10833.

(25) Seal, J. T.; Atkinson, S. J.; Bamborough, P.; Bassil, A.; Chung, C.-W.; Foley, J.; Gordon, L.; Grandi, P.; Gray, J. R. J.; Harrison, L. A.; Kruger, R. G.; Matteo, J. J.; McCabe, M. T.; Messenger, C.; Mitchell, D.; Phillipou, A.; Preston, A.; Prinjha, R. K.; Rianjongdee, F.; Rioja, I.; Taylor, S.; Wall, I. D.; Watson, R. J.; Woolven, J. M.; Wyce, A.; Zhang, X.-P.; Demont, E. H. Fragment-Based Scaffold Hopping: Identification of Potent, Selective, and Highly Soluble Bromo and Extra Terminal Domain (BET) Second Bromodomain (BD2) Inhibitors. J Med Chem 2021, 64 (15), 10772–10805.

(26) Lucas, S. C. C.; Atkinson, S. J.; Chung, C.; Davis, R.; Gordon, L.; Grandi, P.; Gray, J. J. R.; Grimes, T.; Phillipou, A.; Preston, A. G.; Prinjha, R. K.; Rioja, I.; Taylor, S.; Tomkinson, N. C. O.; Wall, I.; Watson, R. J.; Woolven, J.; Demont, E. H. Optimization of a Series of 2,3-Dihydrobenzofurans as Highly Potent, Second Bromodomain (BD2)-Selective, Bromo and Extra-Terminal Domain (BET) Inhibitors. J. Med. Chem. 2021, 64 (15), 10711–10741.

(27) Harrison, L. A.; Atkinson, S. J.; Bassil, A.; Chung, C.-W.; Grandi, P.; Gray, J. R. J.; Levernier, E.; Lewis, A.; Lugo, D.; Messenger, C.; Michon, A.-M.; Mitchell, D. J.; Preston, A.; Prinjha, R. K.; Rioja, I.; Seal, J. T.; Taylor, S.; Wall, I. D.; Watson, R. J.; Woolven, J. M.; Demont, E. H. Identification of a Series of N-Methylpyridine-2-Carboxamides as Potent and Selective Inhibitors of the Second Bromodomain (BD2) of the Bromo and Extra Terminal Domain (BET) Proteins. J Med Chem 2021, 64 (15), 10742–10771.

(28) Gacias, M.; Gerona-Navarro, G.; Plotnikov, A. N.; Zhang, G.; Zeng, L.; Kaur, J.; Moy, G.; Rusinova, E.; Rodriguez, Y.; Matikainen, B.; Vincek, A.; Joshua, J.; Casaccia, P.; Zhou, M.-M. Selective Chemical Modulation of Gene Transcription Favors Oligodendrocyte Lineage Progression. Chem Biol 2014, 21 (7), 841–854.

(29) Cheung, K.; Lu, G.; Sharma, R.; Vincek, A.; Zhang, R.; Plotnikov, A. N.; Zhang, F.; Zhang, Q.; Ju, Y.; Hu, Y.; Zhao, L.; Han, X.; Meslamani, J.; Xu, F.; Jaganathan, A.; Shen, T.; Zhu, H.; Rusinova, E.; Zeng, L.; Zhou, J.; Yang, J.; Peng, L.; Ohlmeyer, M.; Walsh, M. J.; Zhang, D. Y.; Xiong, H.; Zhou, M.-M. BET N-Terminal Bromodomain Inhibition Selectively Blocks Th17 Cell Differentiation and Ameliorates Colitis in Mice. Proc Natl Acad Sci U S A 2017, 114 (11), 2952–2957.

(30) Gilan, O.; Rioja, I.; Knezevic, K.; Bell, M. J.; Yeung, M. M.; Harker, N. R.; Lam, E. Y. N.; Chung, C.; Bamborough, P.; Petretich, M.; Urh, M.; Atkinson, S. J.; Bassil, A. K.; Roberts, E. J.; Vassiliadis, D.; Burr, M. L.; Preston, A. G. S.; Wellaway, C.; Werner, T.; Gray, J. R.; Michon, A.-M.; Gobbetti, T.; Kumar, V.; Soden, P. E.; Haynes, A.; Vappiani, J.; Tough, D. F.; Taylor, S.; Dawson, S.-J.; Bantscheff, M.; Lindon, M.; Drewes, G.; Demont, E. H.; Daniels, D. L.; Grandi, P.; Prinjha, R. K.; Dawson, M. A. Selective Targeting of BD1 and BD2 of the BET Proteins in Cancer and Immunoinflammation. Science 2020, 368 (6489), 387–394.

(31) McLure, K. G.; Gesner, E. M.; Tsujikawa, L.; Kharenko, O. A.; Attwell, S.; Campeau, E.; Wasiak, S.; Stein, A.; White, A.; Fontano, E.; Suto, R. K.; Wong, N. C. W.; Wagner, G. S.; Hansen, H. C.; Young, P. R. RVX-208, an Inducer of ApoA-I in Humans, Is a BET Bromodomain Antagonist. PLoS One 2013, 8 (12), e83190.

(32) Law, R. P.; Atkinson, S. J.; Bamborough, P.; Chung, C.; Demont, E. H.; Gordon, L. J.; Lindon, M.; Prinjha, R. K.; Watson, A. J. B.; Hirst, D. J. Discovery of Tetrahydroquinoxalines as Bromodomain and Extra-Terminal Domain (BET) Inhibitors with Selectivity for the Second Bromodomain. Journal of Medicinal Chemistry 2018, 61(10):4317–4334.

(33) Faivre, E. J.; McDaniel, K. F.; Albert, D. H.; Mantena, S. R.; Plotnik, J. P.; Wilcox, D.; Zhang, L.; Bui, M. H.; Sheppard, G. S.; Wang, L.; Sehgal, V.; Lin, X.; Huang, X.; Lu, X.; Uziel, T.; Hessler, P.; Lam, L. T.; Bellin, R. J.; Mehta, G.; Fidanze, S.; Pratt, J. K.; Liu, D.; Hasvold, L. A.; Sun, C.; Panchal, S. C.; Nicolette, J. J.; Fossey, S. L.; Park, C. H.; Longenecker, K.; Bigelow, L.; Torrent, M.; Rosenberg, S. H.; Kati, W. M.; Shen, Y. Selective Inhibition of the BD2 Bromodomain of BET Proteins in Prostate Cancer. Nature 2020, 578 (7794), 306–310.

(34) Sheppard, G. S.; Wang, L.; Fidanze, S. D.; Hasvold, L. A.; Liu, D.; Pratt, J. K.; Park, C. H.; Longenecker, K.; Qiu, W.; Torrent, M.; Kovar, P. J.; Bui, M.; Faivre, E.; Huang, X.; Lin, X.; Wilcox, D.; Zhang, L.; Shen, Y.; Albert, D. H.; Magoc, T. J.; Rajaraman, G.; Kati, W. M.; McDaniel, K. F. Discovery of N-Ethyl-4-[2-(4-Fluoro-2,6-Dimethyl-Phenoxy)-5-(1-Hydroxy-1-Methyl-Ethyl)Phenyl]-6-Methyl-7-Oxo-1H-Pyrrolo[2,3-c]Pyridine-2-Carboxamide (ABBV-744), a BET Bromodomain Inhibitor with Selectivity for the Second Bromodomain. J. Med. Chem. 2020, 63 (10), 5585–5623.

(35) Preston, A.; Atkinson, S.; Bamborough, P.; Chung, C.-W.; Craggs, P. D.; Gordon, L.; Grandi, P.; Gray, J. R. J.; Jones, E. J.; Lindon, M.; Michon, A.-M.; Mitchell, D. J.; Prinjha, R. K.; Rianjongdee, F.; Rioja, I.; Seal, J.; Taylor, S.; Wall, I.; Watson, R. J.; Woolven, J.; Demont, E. H. Design and Synthesis of a Highly Selective and In Vivo-Capable Inhibitor of the Second Bromodomain of the Bromodomain and Extra Terminal Domain Family of Proteins. J Med Chem 2020, 63 (17), 9070–9092.

(36) Bosc, N.; Muller, C.; Hoffer, L.; Lagorce, D.; Bourg, S.; Derviaux, C.; Gourdel, M.-E.; Rain, J.-C.; Miller, T. W.; Villoutreix, B. O.; Miteva, M. A.; Bonnet, P.; Morelli, X.; Sperandio, O.; Roche, P. Fr-PPIChem: An Academic Compound Library Dedicated to Protein–Protein Interactions. ACS Chem. Biol. 2020, 15(6):1566–1574.

(37) Degorce, F.; Card, A.; Soh, S.; Trinquet, E.; Knapik, G. P.; Xie, B. HTRF: A Technology Tailored for Drug Discovery - a Review of Theoretical Aspects and Recent Applications. Curr Chem Genomics 2009, 3, 22–32.

(38) Chen, Z.; Zhang, H.; Liu, S.; Xie, Y.; Jiang, H.; Lu, W.; Xu, H.; Yue, L.; Zhang, Y.; Ding, H.; Zheng, M.; Yu, K.; Chen, K.; Jiang, H.; Luo, C. Discovery of Novel Trimethoxy-Ring BRD4 Bromodomain Inhibitors: AlphaScreen Assay, Crystallography and Cell-Based Assay †The Authors Declare No Competing Interests. ‡Electronic Supplementary Information (ESI) Available. See DOI: 10.1039/C7md00083a. Medchemcomm 2017, 8 (6), 1322–1331.

(39) Wang, K.; Kim, D.; Dömling, A. Cyanoacetamide MCR (III): Three-Component Gewald Reactions Revisited. J. Comb. Chem. 2010, 12 (1), 111–118.

(40) Mertz, J. A.; Conery, A. R.; Bryant, B. M.; Sandy, P.; Balasubramanian, S.; Mele, D. A.; Bergeron, L.; Sims, R. J. Targeting MYC Dependence in Cancer by Inhibiting BET Bromodomains. Proc Natl Acad Sci U S A 2011, 108 (40), 16669–16674.

(41) Collignon, A.; Hospital, M. A.; Montersino, C.; Courtier, F.; Charbonnier, A.; Saillard, C.; D’Incan, E.; Mohty, B.; Guille, A.; Adelaïde, J.; Carbuccia, N.; Garnier, S.; Mozziconacci, M. J.; Zemmour, C.; Pakradouni, J.; Restouin, A.; Castellano, R.; Chaffanet, M.; Birnbaum, D.; Collette, Y.; Vey, N. A Chemogenomic Approach to Identify Personalized Therapy for Patients with Relapse or Refractory Acute Myeloid Leukemia: Results of a Prospective Feasibility Study. Blood Cancer J 2020, 10 (6), 64.

(42) Karjalainen, R.; Pemovska, T.; Popa, M.; Liu, M.; Javarappa, K. K.; Majumder, M. M.; Yadav, B.; Tamborero, D.; Tang, J.; Bychkov, D.; Kontro, M.; Parsons, A.; Suvela, M.; Mayoral Safont, M.; Porkka, K.; Aittokallio, T.; Kallioniemi, O.; McCormack, E.; Gjertsen, B. T.; Wennerberg, K.; Knowles, J.; Heckman, C. A. JAK1/2 and BCL2 Inhibitors Synergize to Counteract Bone Marrow Stromal Cell-Induced Protection of AML. Blood 2017, 130 (6), 789–802.

(43) Benyamine, A.; Le Roy, A.; Mamessier, E.; Gertner-Dardenne, J.; Castanier, C.; Orlanducci, F.; Pouyet, L.; Goubard, A.; Collette, Y.; Vey, N.; Scotet, E.; Castellano, R.; Olive, D. BTN3A Molecules Considerably Improve Vγ9Vδ2T Cells-Based Immunotherapy in Acute Myeloid Leukemia. Oncoimmunology 2016, 5 (10), e1146843.

(44) Di Veroli, G. Y.; Fornari, C.; Wang, D.; Mollard, S.; Bramhall, J. L.; Richards, F. M.; Jodrell, D. I. Combenefit: An Interactive Platform for the Analysis and Visualization of Drug Combinations. Bioinformatics 2016, 32 (18), 2866–2868.

(45) Bhadury, J.; Nilsson, L. M.; Muralidharan, S. V.; Green, L. C.; Li, Z.; Gesner, E. M.; Hansen, H. C.; Keller, U. B.; McLure, K. G.; Nilsson, J. A. BET and HDAC Inhibitors Induce Similar Genes and Biological Effects and Synergize to Kill in Myc-Induced Murine Lymphoma. PNAS 2014, 111 (26), E2721–E2730.

(46) Dickson, M. A.; Rathkopf, D. E.; Carvajal, R. D.; Grant, S.; Roberts, J. D.; Reid, J. M.; Ames, M. M.; McGovern, R. M.; Lefkowitz, R. A.; Gonen, M.; Cane, L. M.; Dials, H. J.; Schwartz, G. K. A Phase I Pharmacokinetic Study of Pulse-Dose Vorinostat with Flavopiridol in Solid Tumors. Invest New Drugs 2011, 29 (5), 1004–1012.

(47) Amorim, S.; Stathis, A.; Gleeson, M.; Iyengar, S.; Magarotto, V.; Leleu, X.; Morschhauser, F.; Karlin, L.; Broussais, F.; Rezai, K.; Herait, P.; Kahatt, C.; Lokiec, F.; Salles, G.; Facon, T.; Palumbo, A.; Cunningham, D.; Zucca, E.; Thieblemont, C. Bromodomain Inhibitor OTX015 in Patients with Lymphoma or Multiple Myeloma: A Dose-Escalation, Open-Label, Pharmacokinetic, Phase 1 Study. Lancet Haematol 2016, 3 (4), e196–204.

(48) Berthon, C.; Raffoux, E.; Thomas, X.; Vey, N.; Gomez-Roca, C.; Yee, K.; Taussig, D. C.; Rezai, K.; Roumier, C.; Herait, P.; Kahatt, C.; Quesnel, B.; Michallet, M.; Recher, C.; Lokiec, F.; Preudhomme, C.; Dombret, H. Bromodomain Inhibitor OTX015 in Patients with Acute Leukaemia: A Dose-Escalation, Phase 1 Study. Lancet Haematol 2016, 3 (4), e186–195.

(49) Lewin, J.; Soria, J.-C.; Stathis, A.; Delord, J.-P.; Peters, S.; Awada, A.; Aftimos, P. G.; Bekradda, M.; Rezai, K.; Zeng, Z.; Hussain, A.; Perez, S.; Siu, L. L.; Massard, C. Phase Ib Trial With Birabresib, a Small-Molecule Inhibitor of Bromodomain and Extraterminal Proteins, in Patients With Selected Advanced Solid Tumors. J Clin Oncol 2018, 36 (30), 3007–3014.

(50) Zhao, L.; Okhovat, J.-P.; Hong, E. K.; Kim, Y. H.; Wood, G. S. Preclinical Studies Support Combined Inhibition of BET Family Proteins and Histone Deacetylases as Epigenetic Therapy for Cutaneous T-Cell Lymphoma. Neoplasia 2019, 21 (1), 82–92.

(51) Romagnoli, R.; Prencipe, F.; Oliva, P.; Cacciari, B.; Balzarini, J.; Liekens, S.; Hamel, E.; Brancale, A.; Ferla, S.; Manfredini, S.; Zurlo, M.; Finotti, A.; Gambari, R. Synthesis and Biological Evaluation of New Antitubulin Agents Containing 2-(3⍰,4⍰,5⍰-Trimethoxyanilino)-3,6-Disubstituted-4,5,6,7-Tetrahydrothieno[2,3-c]Pyridine Scaffold. Molecules 2020, 25 (7), 1690.

(52) Cai, M.-G.; Wu, Y.; Chang, J. Synthesis and Biological Evaluation of 2-Arylimino-3-Pyridin-Thiazolineone Derivatives as Antibacterial Agents. Bioorg Med Chem Lett 2016, 26 (10), 2517–2520.

(53) Filippakopoulos, P.; Knapp, S. The Bromodomain Interaction Module. FEBS Letters 2012, 586 (17), 2692–2704.

