## Supplementary_Informations for "CRCM5484: A BET- BDII Selective Compound With Differential Anti-Leukemic Drug Modulation"

### Table of contents

**Figure S1. HTRF evaluation of the FR-PPIChem screening hits.** **S3**
**Figure S2. Chemical structures.** **S4**
**Figure S3. Binding mode comparison between CRCM5484 and 4ZW1 (PDB ID).** **S5**
**Figure S4. Selectivity evaluation of compound 12 (A) and compound 16 (B).** **S5**
**Figure S5. Binding mode study of CRCM5484.** **S7**
**Figure S6. CRCM5484 compared to other BDII selective compounds.** **S8**
**Figure S7. CRCM5484 HPLC (A) and LCMS (B) analysis.** **S11**

**Table S1. BROMOscan evaluation of CRCM5484.** **S6**
**Table S2. Maximal concentration used for the 78 evaluated compounds in the drug sensitivity and resistance profiling.** **S9**
**Table S3. Compounds SMILES, provider ID and CAS number.** **S10**
**Table S4. HTRF experimental conditions.** **S11**
**Table S5. X-ray data collection and refinement statistics.** **S12**

### Supplementary information

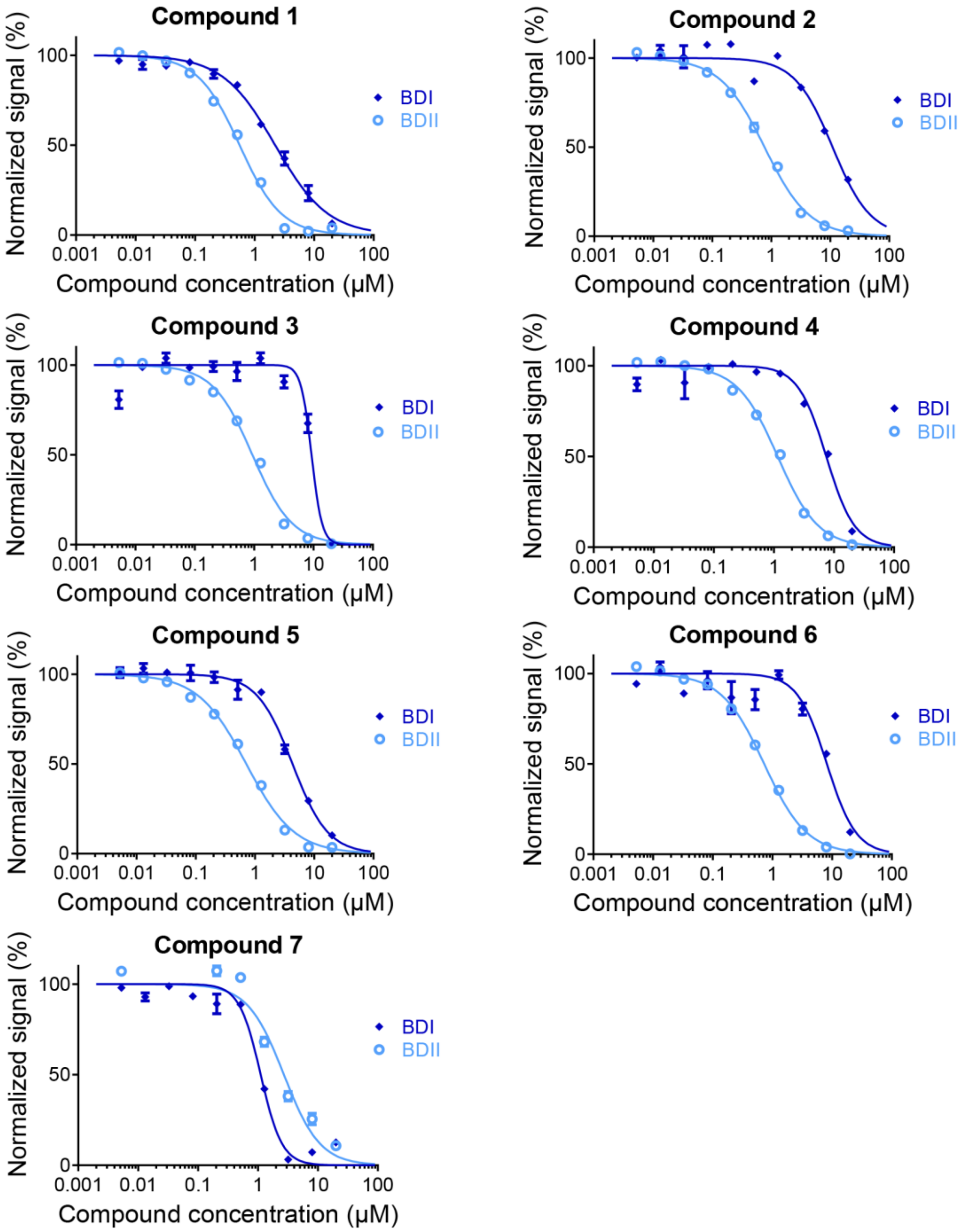


**Supplementary Figure S1. HTRF evaluation of the FR-PPIChem screening hits.** HTRF assay on the BDI (blue lines) and the BDII (cyan lines) domain of BRD3.

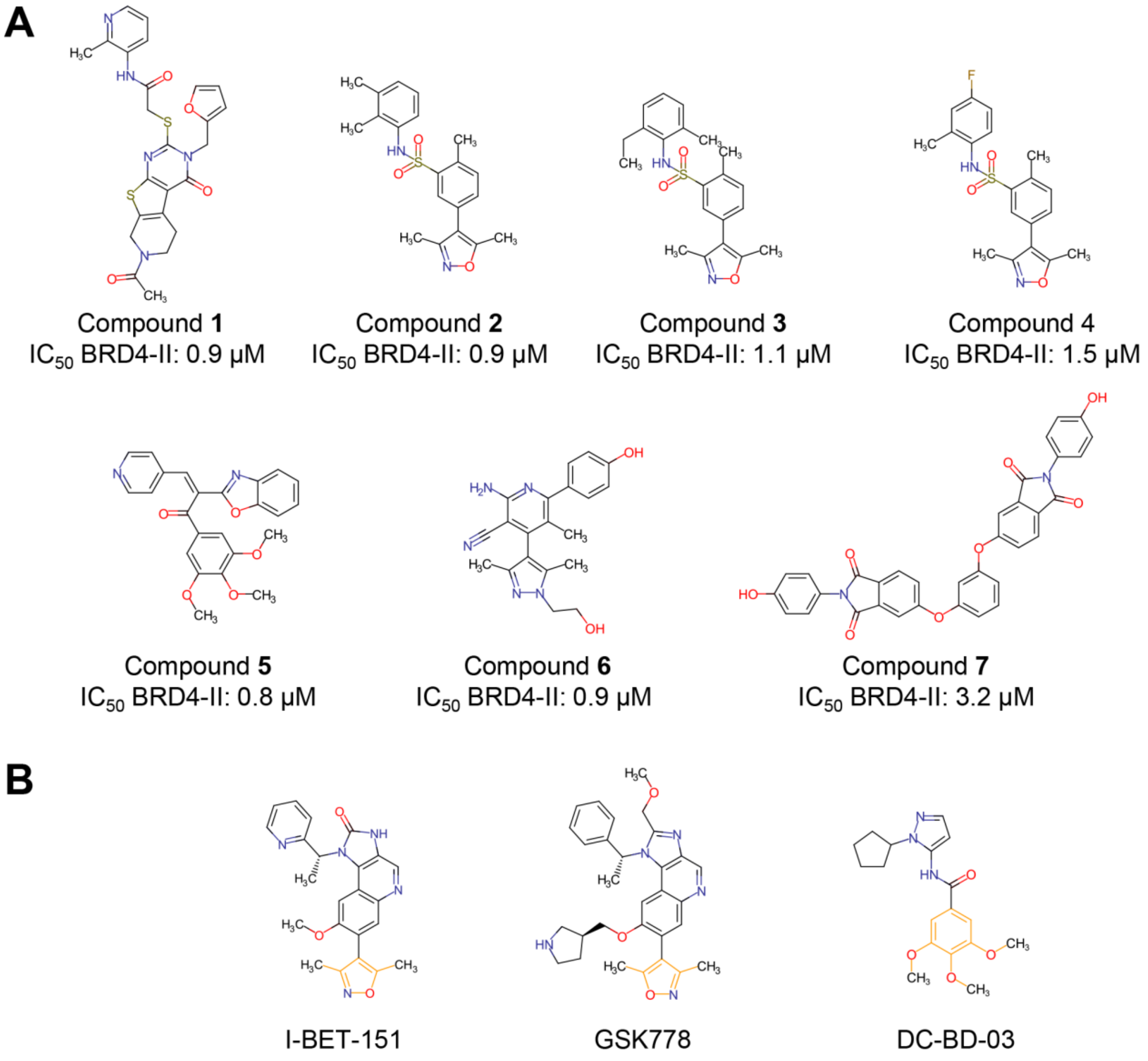


**Figure S2. Chemical structures.** (A) Chemical structure of the seven FR-PPIChem screening hits. (B) Compounds already described with a dimethyl isoxazole or a trimethoxyphenyl core (both colored in orange).

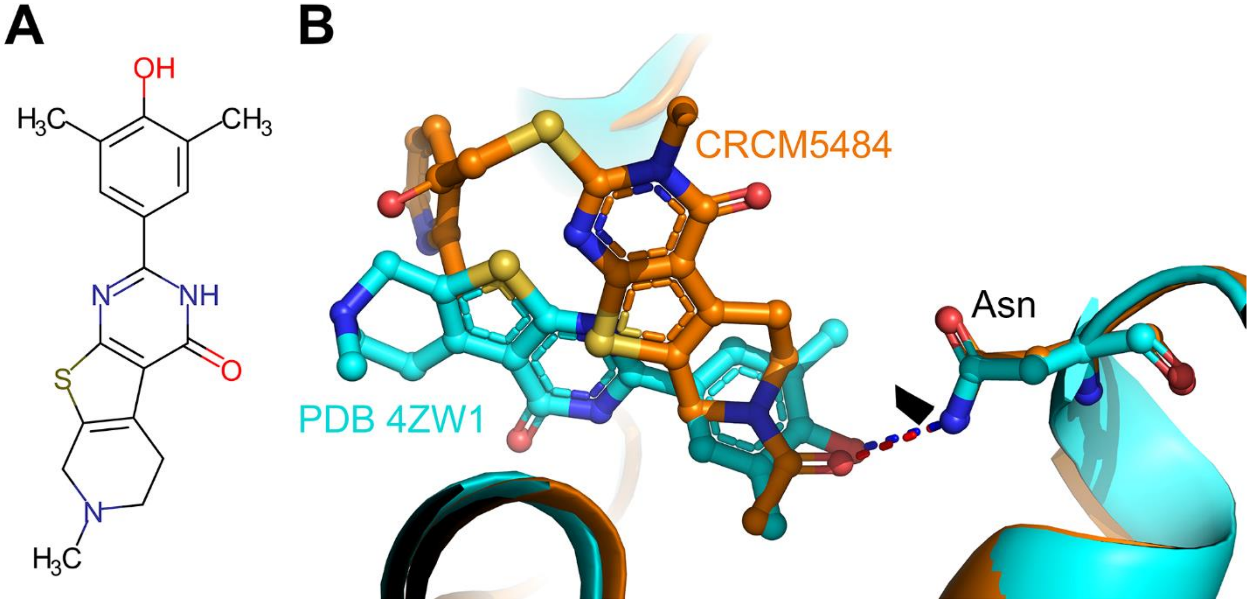


**Figure S3. Binding mode comparison between CRCM5484 and 4ZW1 (PDB ID).** (A) Chemical structure of the 4ZW1 compound. (B) Superposition of the co-crystal structure of CRCM5484)/BRD2-BDII (orange) (PDB: 7Q5O) and 4ZW1 (cyan). The hydrogen bond interactions are shown as red dotted lines for CRCM5484/BRD2-BDII and blue dotted lines for 4ZW1.

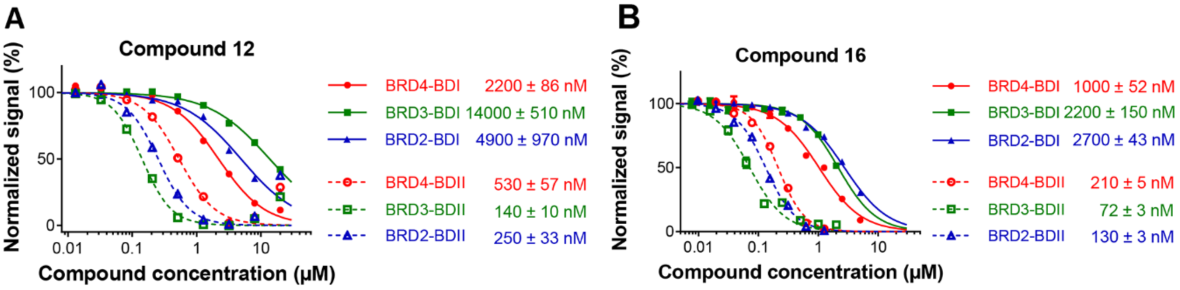


**Figure S4. Selectivity evaluation of compound 12 (A) and compound 16 (B).** HTRF selectivity assay on the BDI (solid lines) and BDII (dashed lines) domain of BRD2 (blue), BRD3 (green) and BRD4 (red).

**Table S1. BROMOscan evaluation of CRCM5484.**

| Target | CRCM5484 |
|---|---|
| Gene Symbol | %Ctrl @ 1 μM |
| ATAD2A | 100 |
| ATAD2B | 90 |
| BAZ2A | 95 |
| BAZ2B | 100 |
| BRD1 | 80 |
| BRD2-I | 92 |
| BRD2-II | 37 |
| BRD3-I | 100 |
| BRD3-II | 36 |
| BRD4-I | 100 |
| BRD4-II | 41 |
| BRD7 | 74 |
| BRD9 | 88 |
| BRDT-I | 96 |
| BRDT-II | 78 |
| BRPF1 | 74 |
| BRPF3 | 100 |
| CECR2 | 54 |
| CREBBP | 85 |
| EP300 | 78 |
| FALZ | 98 |
| GCN5L2 | 99 |
| PBRM1-II | 100 |
| PBRM1-V | 84 |
| PCAF | 100 |
| SMARCA2 | 89 |
| SMARCA4 | 72 |
| TAFI-II | 76 |
| TAF1L-II | 98 |
| TRIM24 | 87 |
| TRIM33 | 52 |
| WDR9 | 91 |

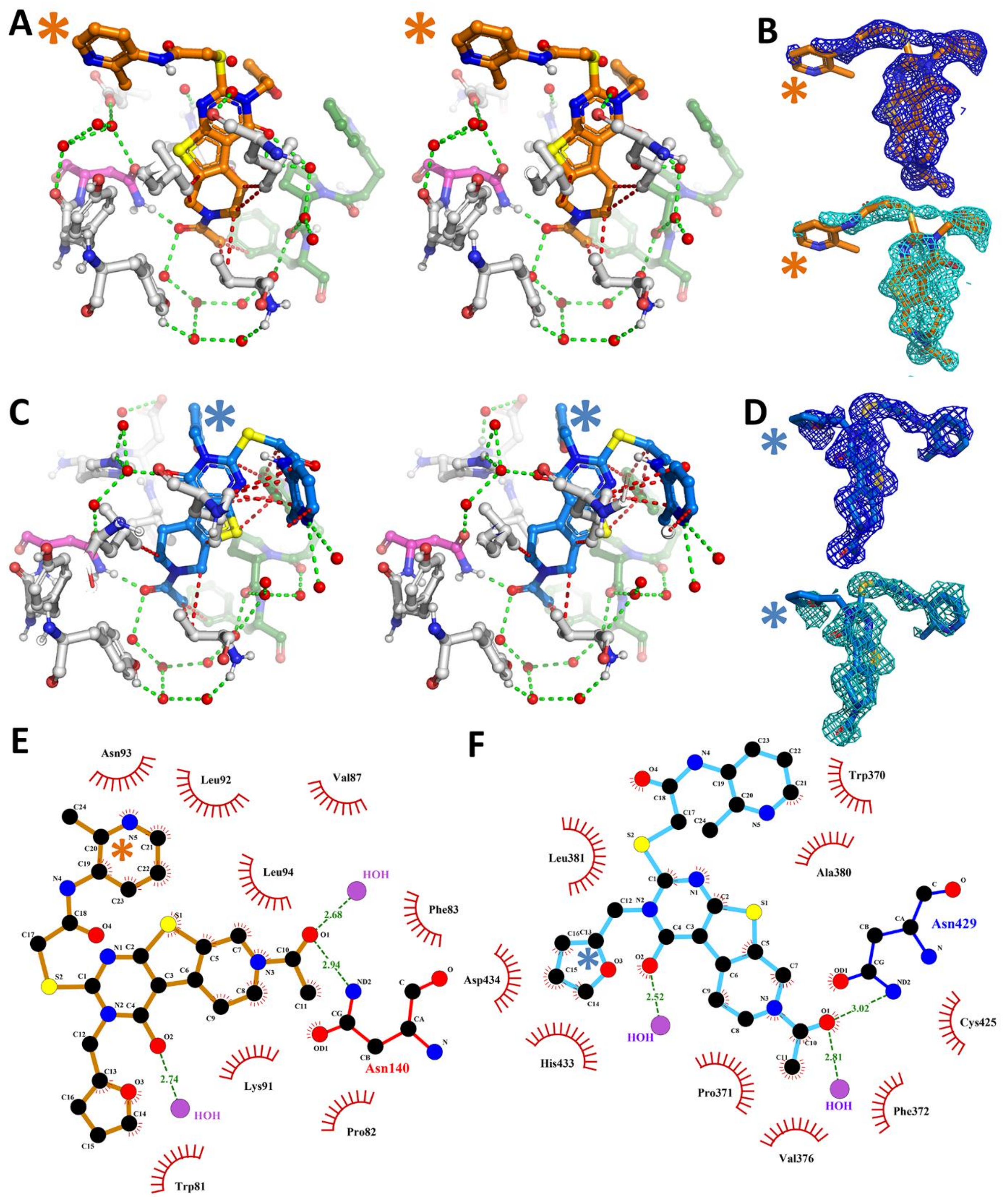


**Figure S5. Binding mode study of CRCM5484.** (A and C) Crossed-eyes stereo views of the co-crystal structures of CRCM5484 with BRD4-BDI (Orange) and BRD2-BDII (Blue) showing the hydrogen bond interactions as green dotted lines, vdW contacts as red dotted lines, the canonical asparagine binding residue as magenta, the WPF shelf as green and the rest of the domain as white balls and sticks. (B and D) 2Fo-Fc (blue) and omitted Fo-Fc map (cyan) of CRCM5484 in the BRD4-BDI (B) and BRD2-BDII (D) X-ray structures. (E and F) 2D Ligplot interaction of the CRCM5484/BRD4-BDI (E) and CRCM5484/BRD2-BDII (F) complexes. The picoline in BRD4-BDI (orange *) and the furan moiety in BRD2-BDII (blue *) showed poor electronic density and their positioning were driven by geometry and residual electron density during refinement.

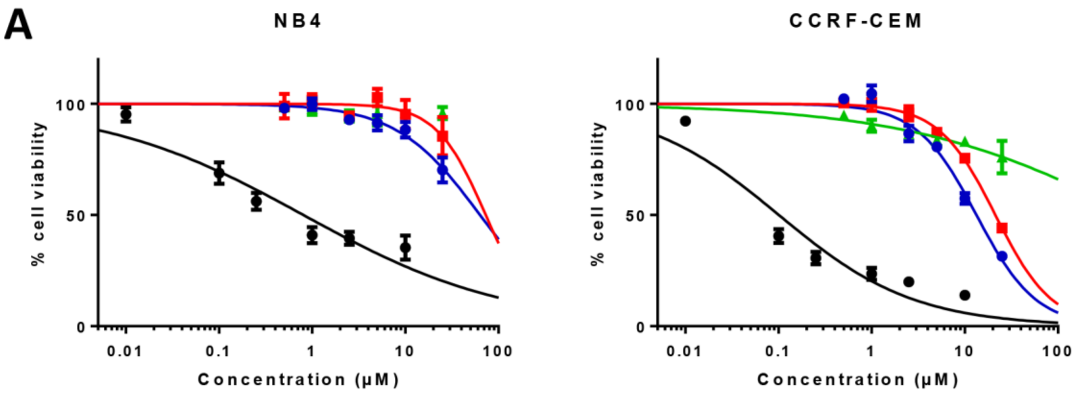


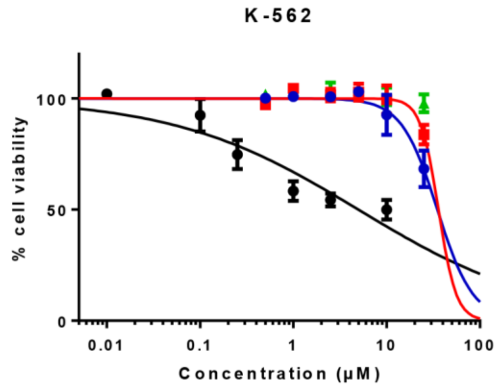


| $EC_{50}$ (µM) | NB4 | CCRF-CEM | K-562 |
|---|---|---|---|
| **OTX-015** | 0.80 | 0.10 | 4.96 |
| **CRCM5484** | >25.00 | 21.28 | >25.00 |
| **GSK046** | >25.00 | >25.00 | >25.00 |
| **RVX-208** | >25.00 | 13.31 | >25.00 |

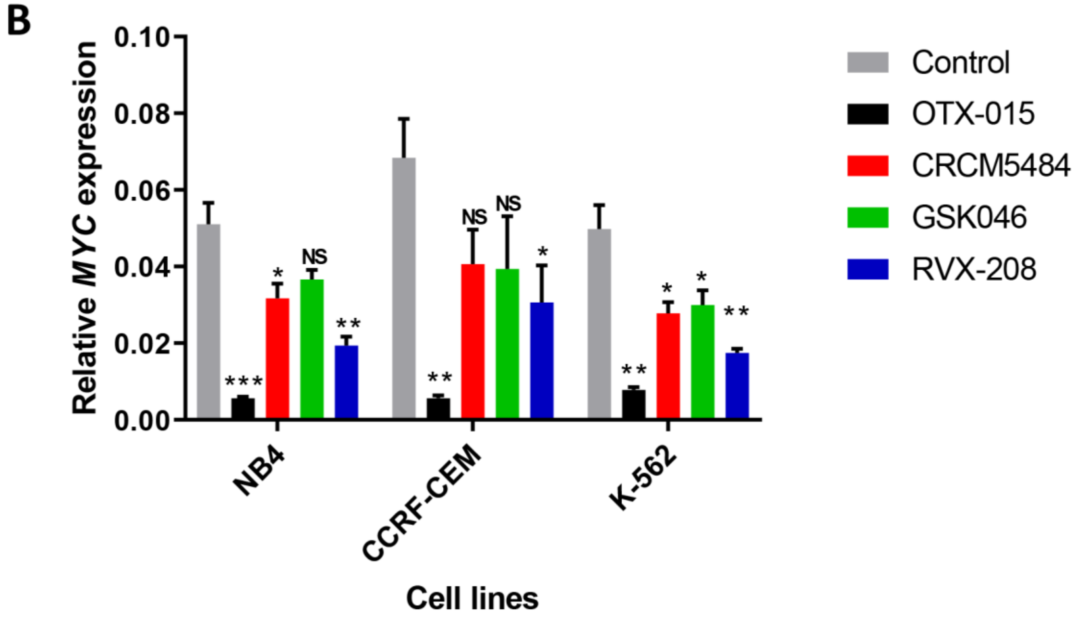


**Figure S6. CRCM5484 compared to other BDII selective compounds.** (A) Cell viability assay on NB4, CCRF-CEM and K-562 leukemic cells performed after 72h incubation with OTX-015 (black), RVX-208 (blue), GSK046 (green) and CRCM5484 (red). (B) Quantification of MYC expression by RT-qPCR following 2h incubation with OTX-015 1µM (black), RVX-208 5µM (blue), GSK046 5µM (green) and CRCM5484 5µM (red). NS: for non-significant.

**Table S2. Maximal concentration used for the 78 evaluated compounds in the drug sensitivity and resistance profiling.** Dose response experiments were done from this condition with three additional 1 log serial dilutions. In these experiments, CRCM5484 was used at the $EC_{40}$ concentration measured for each cell type (12 µM for CCRF-CEM: 6 µM for U-937; 5.7 µM for MOLM-14; 0.6 µM for PDX without BMSC-CM; and 4.3 µM for PDX with BMSC-CM).

| DSRP drugs | Dose-response maximal dose [µM] | DSRP drugs | Dose-response maximal dose [µM] | DSRP drugs | Dose-response maximal dose [µM] |
|---|---|---|---|---|---|
| Luminespid | **0.02** | OTX015 | **20** | Etoposide | **50** |
| Panobinostat | **0.02** | Ponatinib | **20** | Everolimus | **50** |
| Venetoclax | **0.02** | Prima1 | **20** | Gefitinib | **50** |
| Bortezomib | **0.2** | Quizartinib | **20** | GSK126 | **50** |
| Carfilzomib | **0.2** | SGI-1027 | **20** | GSK2879552 | **50** |
| Idarubicin | **0.2** | Sunitinib | **20** | Idasanutlin | **50** |
| Navitoclax | **0.2** | Temsirolimus | **20** | Idelalisib | **50** |
| Birinapant | **2** | Volasertib | **20** | Lapatinib | **50** |
| Cladribine | **2** | Vorinostat | **20** | Masitinib | **50** |
| Daunorubicine | **2** | 17AAG | **50** | Nelarabine | **50** |
| Rigosertib | **2** | AC-4-130 | **50** | Nilotinib | **50** |
| Selinexor | **2** | AG-221 | **50** | Nutlin 3 | **50** |
| Trametinib | **2** | AGI-5198 | **50** | Olaparid | **50** |
| Tretinoin | **2** | Alisertib | **50** | Palbociclib | **50** |
| Vincristine | **2** | Arsenic | **50** | Ruxolitinib | **50** |
| Afatinib | **12** | Azacitidine | **50** | Seliciclib | **50** |
| AraC | **20** | BKM120 | **50** | SGI-110 | **50** |
| AT9283 | **20** | Bleomycine | **50** | SGI-1776 | **50** |
| Clofarabine | **20** | Bosutinib | **50** | Sorafenib | **50** |
| DHAP | **20** | Crenolatinib | **50** | Tipifarnib | **50** |
| Erlotinib | **20** | Cyclopamine | **50** | Topotecan | **50** |
| Gilteritinib | **20** | Dasatinib | **50** | Vermurafenib | **50** |
| Imatinib | **20** | Decitabine | **50** | VX680 | **50** |
| JQ1 | **20** | Docetaxel | **50** | Asparaginase | **100** |
| Midostaurin | **20** | Entinostat | **50** | Hydroxychloroquine | **100** |
| MK2206 | **20** | EPZ5676 | **50** | Metformin | **240000** |

**Table S3. Compounds SMILES, Provider, Provider ID and CAS number.**

| # | SMILES | Provider | Provider ID | CAS number |
|---|---|---|---|---|
| **1** | CC(N(CC1)Cc2c1c(C(N(Cc1ccco1)C(SCC(Nc1c(C)ccc(Cl)c1)=O)=N1)=O)c1s2)=O | ChemDiv | G641-0348 | 1185149-44-3 |
| **2** | Cc1noc(C)c1-c1cc(S(Nc2c(C)c(C)ccc2)(=O)=O)c(C)cc1 | ChemDiv | F176-0440 | 1358716-54-7 |
| **3** | CCc(cccc1C)c1NS(c1c(C)ccc(-c2c(C)onc2C)c1)(=O)=O | ChemDiv | F176-0527 | 1358396-26-5 |
| **4** | Cc1noc(C)c1-c1cc(S(Nc(cc2)c(C)cc2F)(=O)=O)c(C)cc1 | ChemDiv | F176-0574 | 1358583-65-9 |
| **5** | COc1cc(C(/C(/c2nc(cccc3)c3o2)=C/c2ccncc2)=O)cc(OC)c1OC | Vitas-M Lab | STL071346 | 2454175-99-4 |
| **6** | Cc1nn(CCO)c(C)c1-c1c(C)c(nc(N)c1C#N)-c1ccc(O)cc1 | Chembridge | 47081723 | 1351112-74-7 |
| **7** | Oc(cc1)ccc1N(C(c(cc1)c2cc1Oc1cccc(Oc(cc3)cc(C(N4c(cc5)ccc5O)=O)c3C4=O)c1)=O)C2=O | Vitas-M Lab | STK391617 | 295348-36-6 |
| **8** | CC(N(CC1)Cc2c1c(C(N(c1ccccc1)C(SCC(Nc1c(C)ccc(Cl)c1)=O)=N1)=O)c1s2)=O | ChemDiv | G641-0025 | 1217090-90-8 |
| **9** | CC(N(CC1)Cc2c1c(C(N(CC1OCCC1)C(SCC(Nc1c(C)ccc(Cl)c1)=O)=N1)=O)c1s2)=O | ChemDiv | G641-0422 | 1189972-54-0 |
| **10** | CCC(N(CC1)Cc2c1c(C(N(C1CCCCC1)C(N1)=O)=O)c1s2)=O | ChemDiv | V014-6657 | 922356-59-0 |
| **11** | CC(=O)N1CCc2c(C1)sc1nc(SCC(=O)Nc3ccccc3)n(-c3ccccc3)c(=O)c21 | ChemDiv | G641-0030 | 1184977-70-5 |
| **12** | CC(N(CC1)Cc2c1c(C(N(Cc1ccco1)C(SCC(Nc1ccccc1)=O)=N1)=O)c1s2)=O | ChemDiv | G641-0355 | 1215718-74-3 |
| **13** | CC(N(CC1)Cc2c1c(C(N(Cc1ccco1)C(SCC(Nc1cc(OC)ccc1)=O)=N1)=O)c1s2)=O | ChemDiv | G641-0323 | 1189643-36-4 |
| **14** | CC(N(CC1)Cc2c1c(C(N(Cc1ccco1)C(SCC(Nc(cccc1)c1Cl)=O)=N1)=O)c1s2)=O | ChemDiv | G641-0325 | 1215508-33-0 |
| **15** | CC(N(CC1)Cc2c1c(C(N(Cc1ccco1)C(SCC(Nc1cc(C)ccc1)=O)=N1)=O)c1s2)=O | ChemDiv | G641-0345 | 1185063-82-4 |
| **16** | CC(N(CC1)Cc2c1c(C(N(Cc1ccco1)C(SCC(Nc(cccc1)c1OC)=O)=N1)=O)c1s2)=O | ChemDiv | G641-0327 | 1189940-97-3 |
| **17** | CC(N(CC1)Cc2c1c(C(N(Cc1ccco1)C(SCC(Nc(cc1)ccc1OC)=O)=N1)=O)c1s2)=O | ChemDiv | G641-0342 | 1215387-46-4 |
| **18** | CC(N(CC1)Cc2c1c(C(N(Cc1ccco1)C(SCC(Nc(cc1)ccc1F)=O)=N1)=O)c1s2)=O | ChemDiv | G641-0343 | 1217036-58-2 |
| **19** | CC(N(CC1)Cc2c1c(C(N(Cc1ccco1)C(SCC(Nc1c(C)cccc1)=O)=N1)=O)c1s2)=O | ChemDiv | G641-0344 | 1216802-03-7 |
| **20** | CC(N(CC1)Cc2c1c(C(N(Cc1ccco1)C(SCC(Nc1ccc(C)cc1)=O)=N1)=O)c1s2)=O | ChemDiv | G641-0326 | 1215481-48-3 |
| **21** | CC(N(CC1)Cc2c1c(C(N(Cc1ccco1)C(SCC(Nc1cccc(Cl)c1)=O)=N1)=O)c1s2)=O | ChemDiv | G641-0346 | 1185044-25-0 |
| **22** | CCOc(cccc1)c1NC(CSC(N1Cc2ccco2)=Nc(sc(C2)c3CCN2C(C)=O)c3C1=O)=O | ChemDiv | G641-0347 | 1216929-41-7 |
| **23** | CC(N(CC1)Cc2c1c(C(N(Cc1ccco1)C(SCC(Nc(ccc(F)c1)c1F)=O)=N1)=O)c1s2)=O | ChemDiv | G641-0351 | 1216742-87-8 |
| **24** | CC(N(CC1)Cc2c1c(C(N(Cc1ccco1)C(SCC(Nc(cccc1)c1F)=O)=N1)=O)c1s2)=O | ChemDiv | G641-0352 | 1185075-81-3 |
| **25** | CC(N(CC1)Cc2c1c(C(N(Cc1ccco1)C(SCC(Nc1cc(F)c(C)cc1)=O)=N1)=O)c1s2)=O | ChemDiv | G641-0353 | 1189947-24-7 |
| **26** | CC(N(CC1)Cc2c1c(C(N(Cc1ccco1)C(SCC(Nc1cc(SC)ccc1)=O)=N1)=O)c1s2)=O | ChemDiv | G641-0357 | 1189692-93-0 |
| **27** | CC(N(CC1)Cc2c1c(C(N(Cc1ccco1)C(SCC(Nc(cc1)ccc1Cl)=O)=N1)=O)c1s2)=O | ChemDiv | G641-0358 | 1216506-86-3 |
| **28** | CC(N(CC1)Cc2c1c(C(N(Cc1ccco1)C(SCC(Nc(cc(cc1)F)c1F)=O)=N1)=O)c1s2)=O | ChemDiv | G641-0361 | 1189906-77-1 |
| **29** | CC(c1cccc(NC(CSC(N2Cc3ccco3)=Nc(sc(C3)c4CCN3C(C)=O)c4C2=O)=O)c1)=O | ChemDiv | G641-0362 | 1189461-87-7 |
| **30** | CC(c(cc1)ccc1NC(CSC(N1Cc2ccco2)=Nc(sc(C2)c3CCN2C(C)=O)c3C1=O)=O)=O | ChemDiv | G641-0366 | 1189944-52-2 |
| **31** | CCOc(cc1)ccc1NC(CSC(N1Cc2ccco2)=Nc(sc(C2)c3CCN2C(C)=O)c3C1=O)=O | ChemDiv | G641-0368 | 1189420-68-5 |
| **32** | CC(N(CC1)Cc2c1c(C(N(Cc1ccco1)C(SCC(Nc(cc1)cc(Cl)c1F)=O)=N1)=O)c1s2)=O | ChemDiv | G641-0373 | 1189437-15-7 |
| **33** | CC(N(CC1)Cc2c1c(C(N(Cc1ccco1)C(SCC(Nc(cc1)cc(F)c1F)=O)=N1)=O)c1s2)=O | ChemDiv | G641-0374 | 1189484-35-2 |
| **34** | CC(N(CC1)Cc2c1c(C(N(Cc1ccco1)C(SCC(Nc1c(C)nccc1)=O)=N1)=O)c1s2)=O | Synthesized | - | - |

**Figure S7. CRCM5484 HPLC (A) and LCMS (B) analysis.**

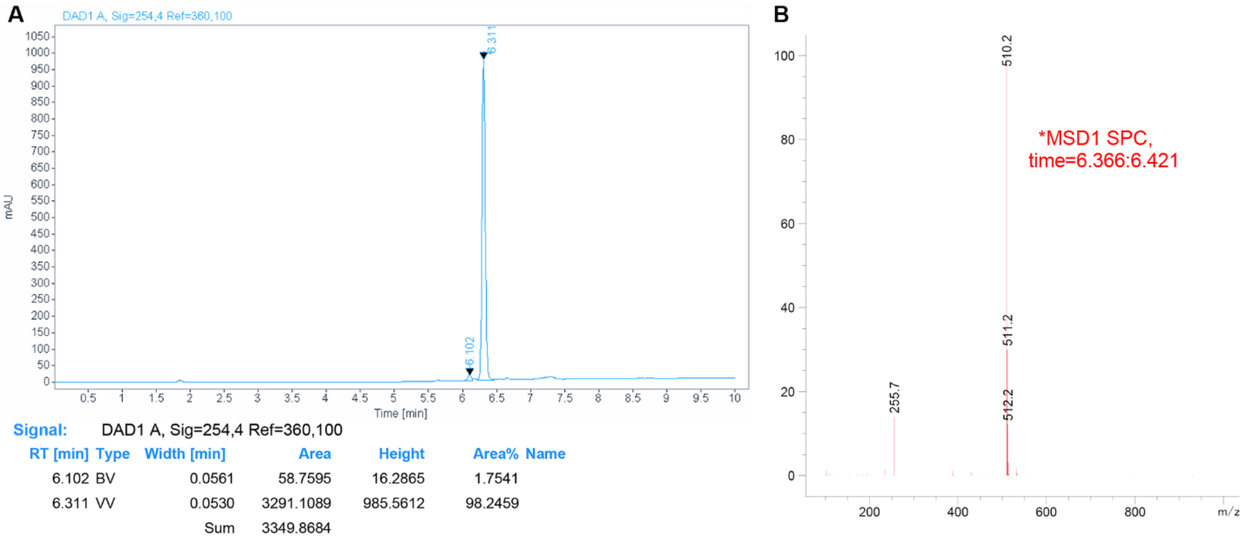


**Table S4. HTRF experimental conditions.**

| HTRF reagents | Details and concentrations [nM] | | Provider |
|---|---|---|---|
| BRDs | GST-BRD4-BDI | GST-BRD4-BDII | Internally produced |
| | 5 | 10 | |
| Donor | MAB Anti GST-EU cryptate | | Cisbio |
| | 0.5 | 0.5 | |
| Acceptor | Streptavidin-d2 (BRD4-BDI) | Streptavidin-XL665 (BRD4-BDII) | Cisbio |
| | 12.5 | 18.75 | |
| Peptide | SGRG-K(Ac)-GG-K(Ac)-GLG-K(Ac)-GGAK(Ac)-RHRKVGG-K(Biotin) >95% Purity | | Genicbio |
| | 15 | 100 | |

**Table S5. X-ray data collection and refinement statistics.**

| | CRCM5484/BRD4-BDI | CRCM5484/BRD2-BDII |
|---|---|---|
| **PDB ID** | 7Q3F | 7Q5O |
| **Crystal parameters** | | |
| Space group | P 21 21 21 | P 21 21 2 |
| Unit-cell dimension | | |
| a, b, c (Å) | 43.21 48.72 61.58 | 52.43 72.49 32.04 |
| α, β, ϒ (°) | 90 90 90 | 90 90 90 |
| **Data collection** | | |
| Beamline | ESRF - ID30A-1 | ESRF - ID30A-1 |
| Total reflections | 148548 (14126) | 75416 (6970) |
| Unique reflection | 40010 (3926) | 19190 (1857) |
| Completeness (%) | 98.07 (88.51) | 98.26 (96.43) |
| Mean I/sigma(I) | 8.6 (0.8) | 4.6 (1.5) |
| R-meas (%) | 50 (180) | 17 (107) |
| CC1/2 | 99.8 (23.3) | 95.5 (4.4) |
| **Structure Refinement** | | |
| Resolution range (Å) | 38.207 - 1.210 (1.253 - 1.21) | 29.82 - 1.52 (1.574 - 1.52) |
| Reflections used in refinement | 39573 (3521) | 19121 (1801) |
| Reflections used for R-free | 1969 (185) | 876 (76) |
| R-work | 0.2056 (0.4408) | 0.2162 (0.3395) |
| R-free | 0.2242 (0.4134) | 0.2464 (0.2990) |
| Number of non-hydrogen atoms | 1246 | 1146 |
| Macromolecules | 1068 | 940 |
| Ligand | 35 | 47 |
| Solvent | 143 | 159 |
| Average B-factor | 28.1 | 14.90 |
| Macromolecules | 26.4 | 12.40 |
| Ligand | 48.1 | 28.00 |
| Solvent | 35.8 | 25.60 |
| Ramachandran plot (%) | | |
| Favored regions | 99.19 | 99.07 |
| Allowed regions | 0.81 | 0.93 |
| Outlier regions | 0.00 | 0 |
| RMS deviation | | |
| Bond (Å) | 0.0102 | 0.0024 |
| Angle (°) | 1.18 | 0.60 |
