## Supplementary material for "CRCM5484: A BET- BDII Selective Compound With Differential Anti-Leukemic Drug Modulation": Providers_Data: V014-6657.PDF

| | | |
|---|---|---|
| Sample: 15 | Date: 21-Jun-2008 | Page 1. |
| File: Ar24505_05 | Time: 01:44:22 | AMRI code: ALB-H10714879 |
| Vial: E/1 | Description: 20579700 | Vial label: 226752-1 |

### ELSD

max. intensity: 2.7E4

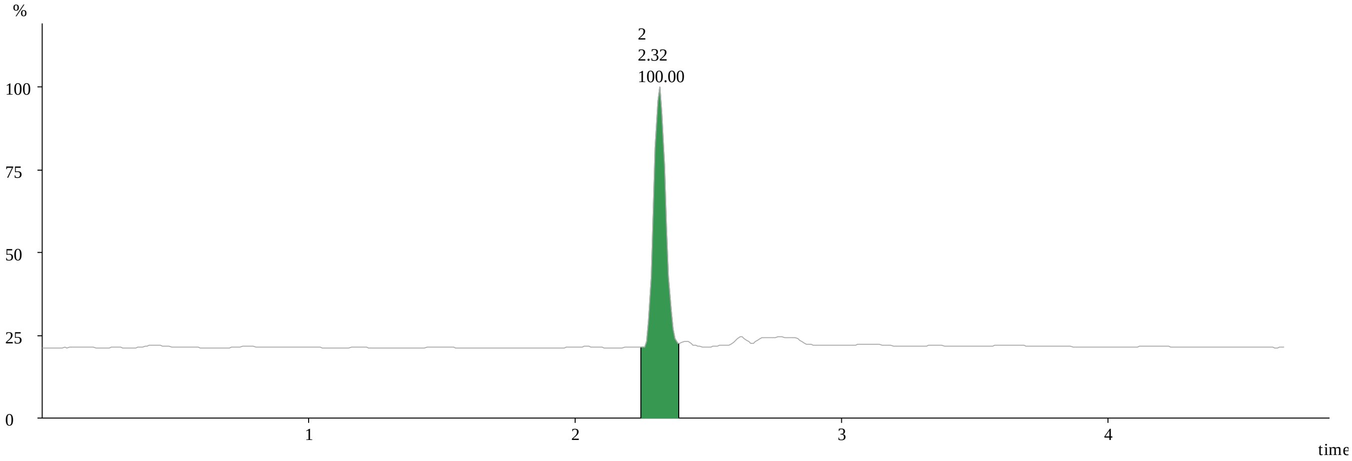


| Peak_ID | Peak | Area | Area% | Height | Time | Mass Found |
|---|---|---|---|---|---|---|
| 2 | 2.25 2.39 | 1.E3 | 100 | 2.E4 | 2.32 | 361.15 |

### DAD: 220

max. intensity: 2.8E5

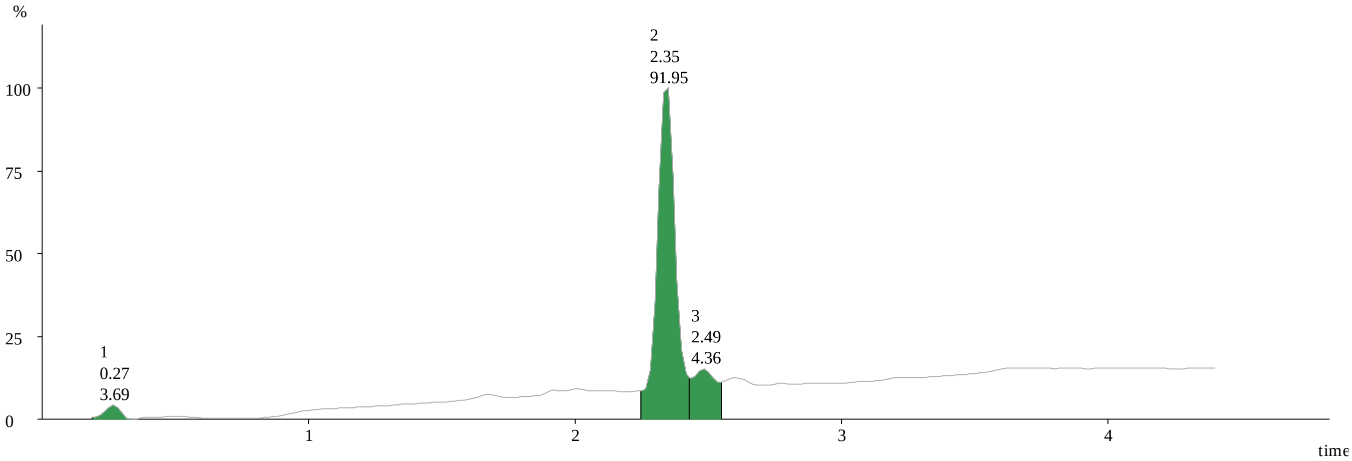


| Peak_ID | Peak | Area | Area% | Height | Time | Mass Found |
|---|---|---|---|---|---|---|
| 1 | 0.19 0.34 | 7.E2 | 3.69 | 1.E4 | 0.27 | 361.15 |
| 2 | 2.25 2.43 | 2.E4 | 91.95 | 3.E5 | 2.35 | 361.15 |
| 3 | 2.43 2.55 | 8.E2 | 4.36 | 1.E4 | 2.49 | 361.15 |

## MS ES+ :723.3+379.15+362.15

max. intensity: 8.8E1

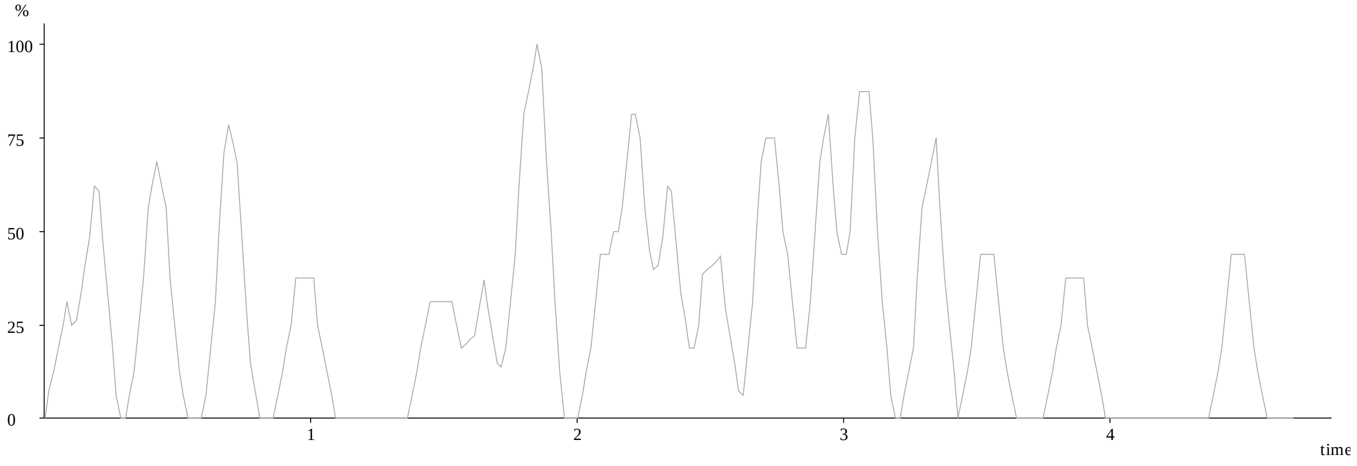


### MS ES+ :TIC

max. intensity: 8.1E3

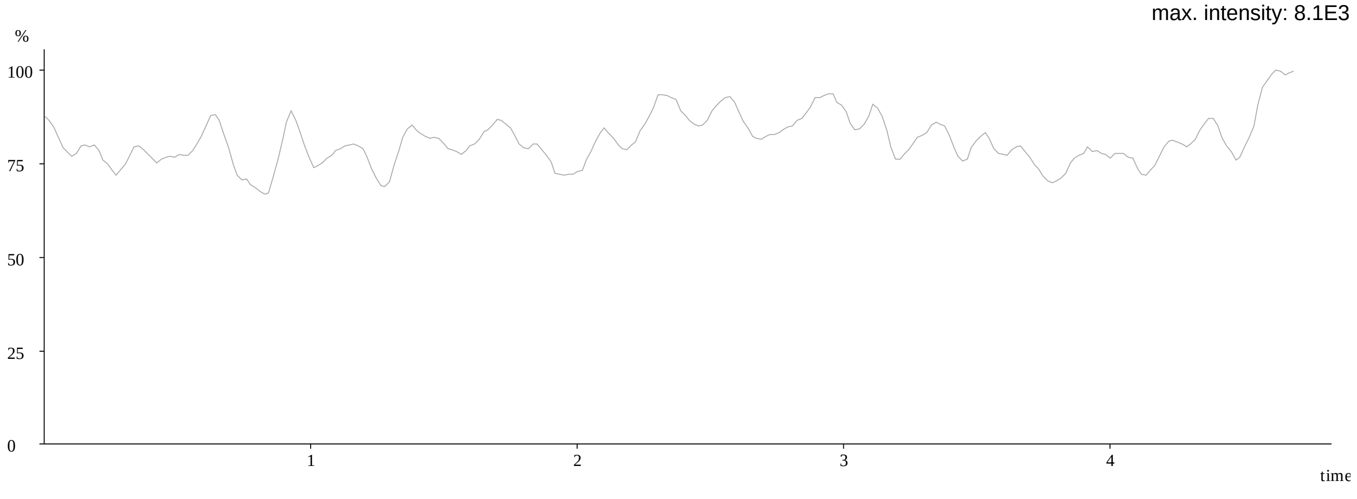

## MS: ES+

Combine (16:18-(8:9+23:25))

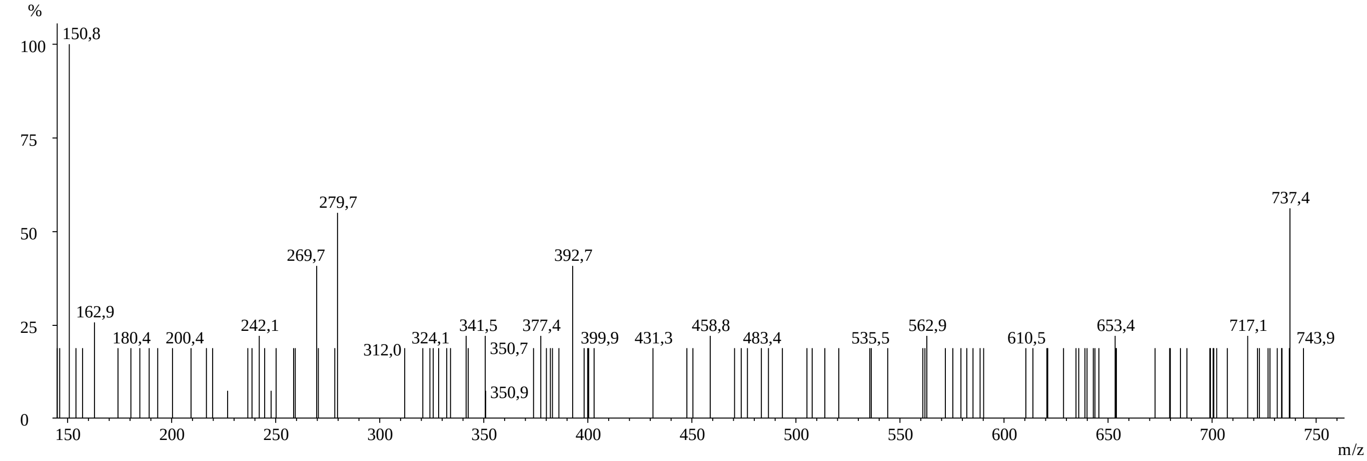


| Peak_ID | Compound | Time | Mass found |
|---|---|---|---|
| 1 | Tentative | 0.27 | 361.1500 |

## MS: ES+

Combine (138:140-(131:132+146:147))

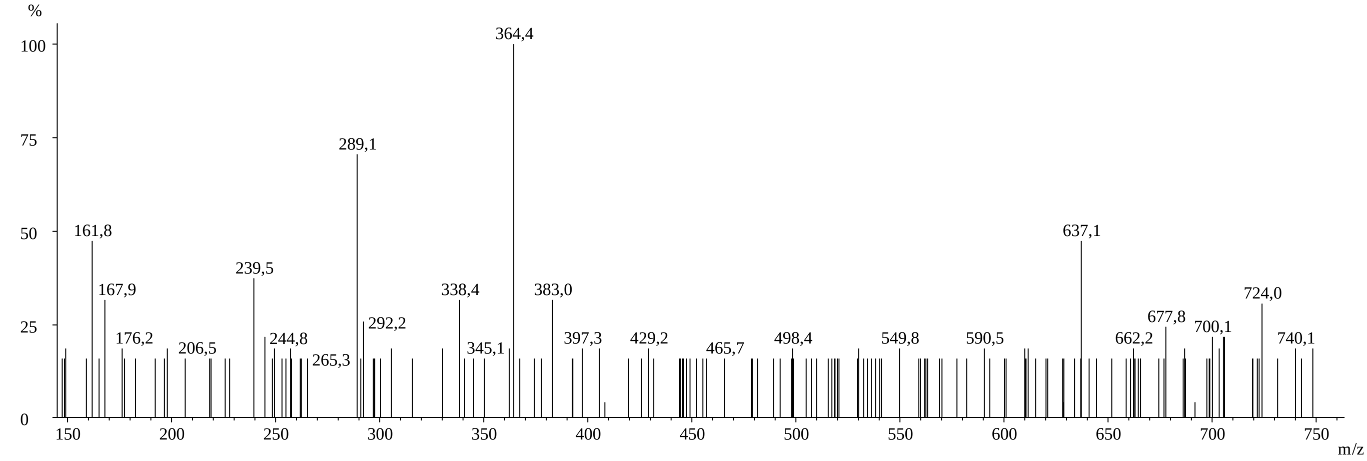


| Peak_ID | Compound | Time | Mass found |
|---|---|---|---|
| 2 | Tentative | 2.32 | 361.1500 |

## MS: ES+

Combine (148:150-(142:143+155:157))

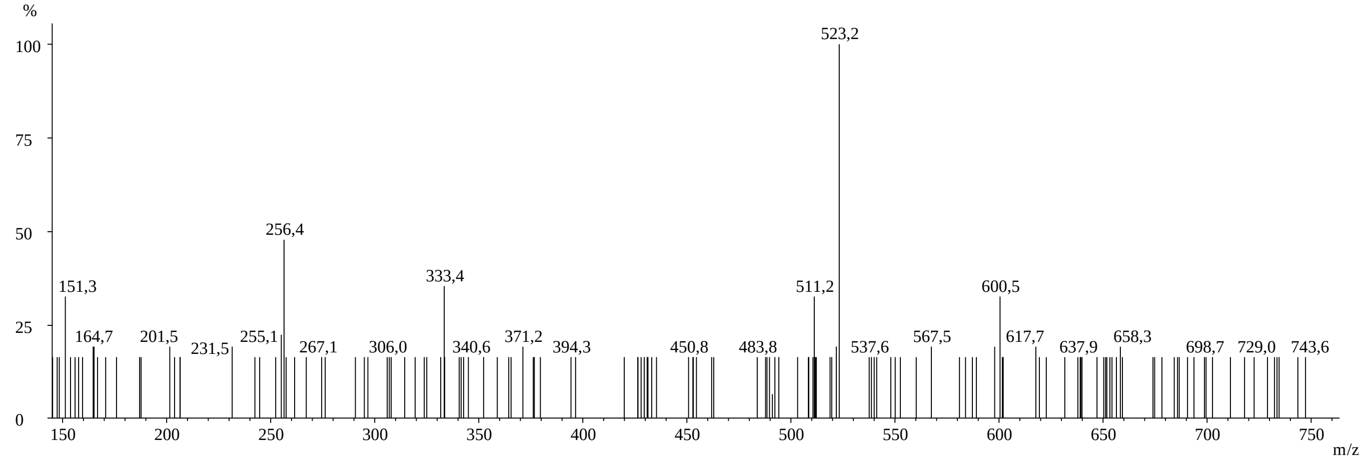


| Peak_ID | Compound | Time | Mass found |
|---|---|---|---|
| 3 | Tentative | 2.49 | 361.1500 |
