## Supplementary figures and images for "CRCM5484: A BET- BDII Selective Compound With Differential Anti-Leukemic Drug Modulation"

### F176-0440.JPG

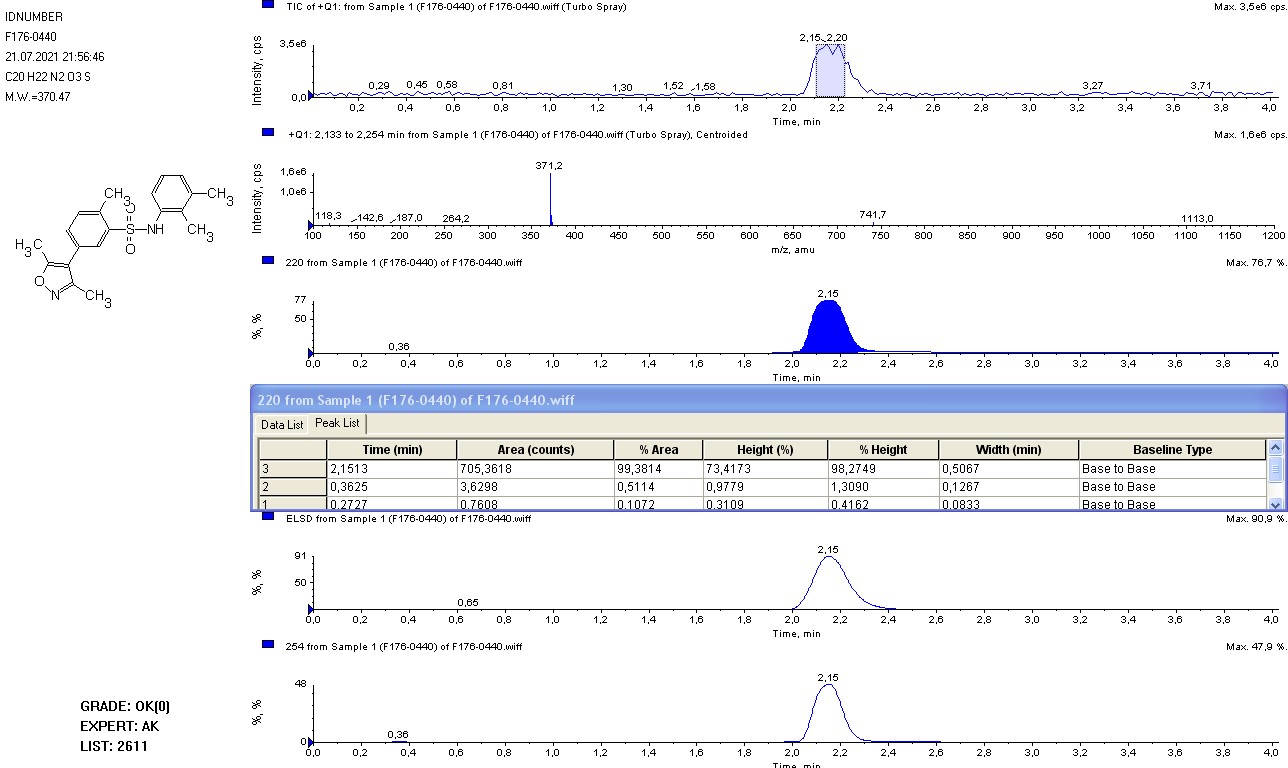

### STK391617.tif

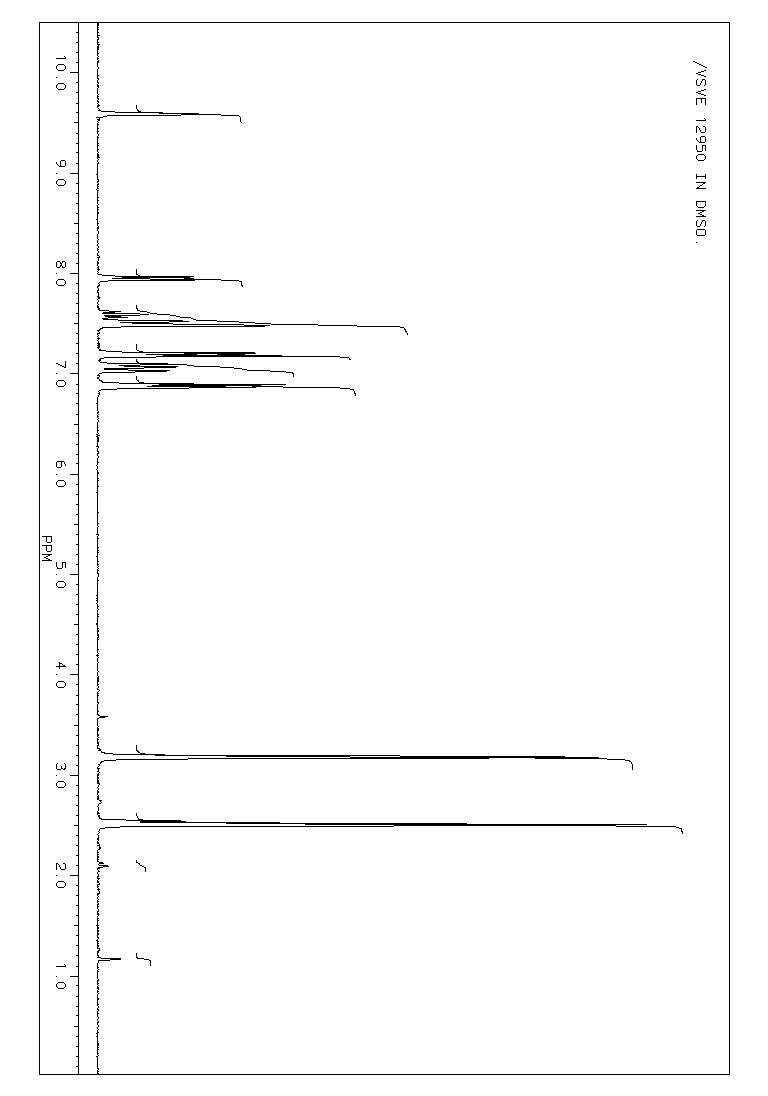

### STL071346.pdf

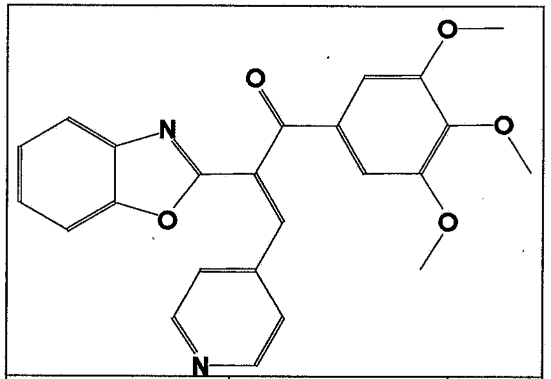

| EX00144134 | C24 H20 N2 O5 | 416.43 |
|---|---|---|
| P2001S-212564 | | |

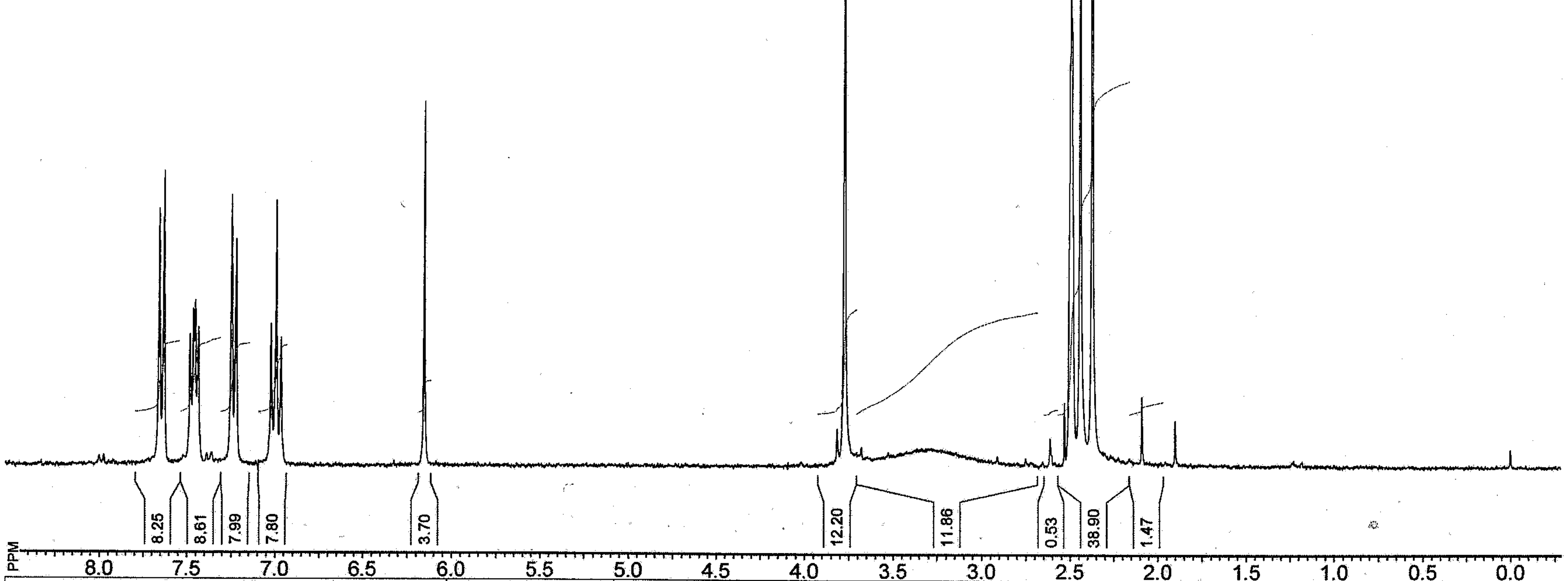


| | | | | |
|---|---|---|---|---|
| **File name: EX00144134** | Owner: | SF: 299.9450 MHz | NS: -1 | SI: 16384, TD: 32768 |
| Date: 30-Dec-1899 | Solvent: | SW: 5099 | TE: 0 | 1137 in DMSO-D6 |
